# Deep learning enables fast and dense single-molecule localization with high accuracy

**DOI:** 10.1101/2020.10.26.355164

**Authors:** Artur Speiser, Lucas-Raphael Müller, Ulf Matti, Christopher J. Obara, Wesley R. Legant, Anna Kreshuk, Jakob H. Macke, Jonas Ries, Srinivas C. Turaga

**Author notes:** These authors contributed equally: Artur Speiser, Lucas-Raphael Müller. For correspondence, Jakob H. Macke, Jonas Ries, Srinivas C. Turaga. These three authors contributed equally.

## Abstract

Single-molecule localization microscopy (SMLM) has had remarkable success in imaging cellular structures with nanometer resolution, but the need for activating only single isolated emitters limits imaging speed and labeling density. Here, we overcome this major limitation using deep learning. We developed DECODE, a computational tool that can localize single emitters at high density in 3D with highest accuracy for a large range of imaging modalities and conditions. In a public software benchmark competition, it outperformed all other fitters on 12 out of 12 data-sets when comparing both detection accuracy and localization error, often by a substantial margin. DECODE allowed us to take live-cell SMLM data with reduced light exposure in just 3 seconds and to image microtubules at ultra-high labeling density. Packaged for simple installation and use, DECODE will enable many labs to reduce imaging times and increase localization density in SMLM.

## Introduction

Single-molecule localization microscopy (SMLM) (e.g. PALM^1^ and STORM^2^) has become an invaluable super-resolution method for biology, as it can resolve cellular structures with nanometer precision. It is based on acquiring a large number of camera frames, in each of which only a tiny fraction of the emitters are activated into a bright ‘on’ state, so that their images do not overlap. This allows precise localization of the emitter coordinates by fitting a model of the Point Spread Function (PSF). A super-resolution image is then reconstructed from these coordinates. This principle of SMLM is at the same time one of its main limitations: the need for sparse activation leads to long acquisition times. This results in low throughput, poor time resolution when imaging dynamic processes, low labeling densities and a reduced choice of fluorophores. Additionally, long acquisition times in combination with high excitation laser intensities needed for single-molecule imaging cause strong photo toxicity in live-cell SMLM.

All of these limitations can be mitigated by activating emitters at a higher density. In this ‘multi-emitter’ setting, PSFs are no longer well-separated but may overlap, making both the detection of multiple nearby emitters and their accurate localization computationally challenging. This is not adequately addressed by existing algorithms: Current ‘multi-emitter’ fitting algorithms work reasonably well on two-dimensional samples where all emitters have the same z-coordinate and thus produce identical PSFs. These algorithms, however, have had limited success for realistic three-dimensional biological structures. In a software competition that benchmarked SMLM algorithms using realistic computer-generated data, simple single-emitter fitters outperformed dedicated high-density fitters on three-dimensional samples even in the high density regime^3^.

Deep learning is revolutionizing biological image analysis^4–6^. For SMLM, deep learning holds promise to extract emitter coordinates and additional parameters under conditions and densities too complex for traditional fitters. With enough training data, deep networks are flexible function approximators which can be trained to recognize patterns in the image and thus transform images directly into predicted configurations of emitters, even for challenging high densities of emitters. While ground-truth data to train the neural network is typically not available, synthetic training data can be generated by numerically simulating the imaging process^7,8^. Convolutional neural networks (CNNs, a class of deep networks suitable for image data) have recently been used to extract parameters describing single isolated emitters such as color, emitter orientation, z-coordinate, background or aberrations^9–12^ and to design optimized PSFs^13^. Two recent studies (DeepSTORM3D^13^ and DeepLoco^14^) used CNNs for extracting emitter coordinates, and outperformed traditional single-emitter fitting algorithms at densities higher than the single-molecule regime. These studies illustrate the potential of deep learning for SMLM, however they have only been demonstrated either for exotic engineered point spread functions or on simulated data.

Here we present the DECODE (DEep COntext DEpendent) method for deep-learning based single-molecule localization that achieves high accuracy across a wide range of emitter densities and brightnesses. DECODE uses a novel deep network output representation, architecture, and cost function, which enable simultaneous detection and sub-pixel localization of single emitters. Uniquely, DECODE is able to predict both the probability of detection and the uncertainty of localization for each emitter. As the timing and duration of emitter activations are stochastic, they regularly persist over several imaging frames. The DECODE architecture can integrate information across neighboring frames (‘temporal context’), which improves emitter detection and localization.

In the public SMLM challenge^3^, DECODE outperformed all existing methods on 12 out of 12 datasets. Compared to previous deep learning based high-density fitters^13^, DECODE is 10x faster and up to 2x more accurate, and it can be applied to a wide range of PSFs. We demonstrate on biological structures that DECODE allows for 6-fold higher labeling densities or 10-fold faster imaging compared to imaging in the single emitter regime, and thus enables fast live-cell SMLM with reduced light exposure. We show the versatility of DECODE by re-analyzing a published Lattice Light-Sheet PAINT data set^15^ for which we could substantially improve fluorophore detection and localization accuracy. DECODE is packaged for simple use and can be easily trained and used by non-expert users, without having to design new network architectures. Thus, it will enable the entire community to overcome the need of sparse activation as one of the main bottlenecks in SMLM.

## Results

### DECODE network

DECODE introduces a new output representation and architecture for detecting and localizing emitters. For each image frame it predicts multiple channels with the same dimensions as the input image (Fig. 1a). The first two channels indicate the probability *p* that an emitter exists near that pixel, as well as its brightness *N* (number of photons emitted by the emitter in the frame). The next three channels describe the coordinates of the emitter with respect to the center of the pixel, Δ*xyz* = [Δ*x*, Δ*y*, Δ*z*]. An additional channel predicts the background intensity *B* in each pixel.

**Figure 1.**
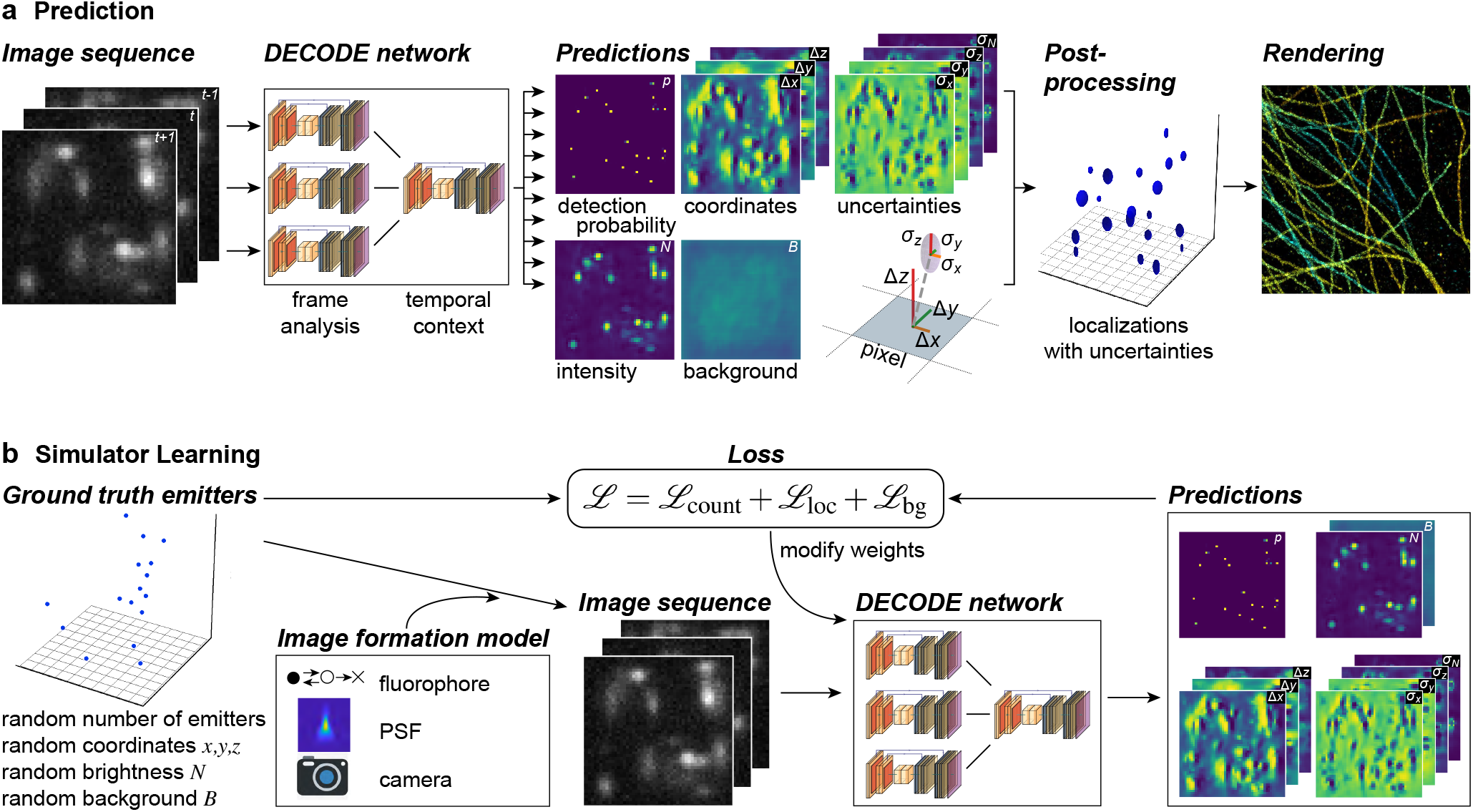
DECODE for high-density single molecule localization. **a) DECODE architecture.** The DECODE network uses information from multiple frames to predict detection probabilities, coordinates and uncertainty estimates. The *frame analysis module* with a multi-scale ‘U-Net’ architecture^16^ extracts informative features from each frame. These features are integrated by the *temporal context module* which produces 9 output maps: a map of emitter detection probabilities *p*, a map predicting the brightness of the corresponding detected emitter *N*, three maps of the three spatial coordinates of the detected emitter Δ*x*, Δ*y*, Δ*z* (relative to the to the center of the detected pixel) and four maps of the associated uncertainties (standard deviations) *σ N, σx, σy, σz*. In addition, we optionally predict a map with the background intensity *B* in each pixel. **b) Training DECODE.** The DECODE network is trained by simulator-learning: Ground truth (GT) emitter coordinates are generated randomly and a forward-model of the image formation process is used to simulate synthetic images. These simulated images are passed through the DECODE network. The loss quantifies the probability that the GT explains the output predictions. This probability is maximized during training.

This architecture overcomes limitations of current deep-learning^13,14^ and non-deep learning based high density approaches in three ways: First, DECODE predictions scale only with the number of imaged pixels (not super-resolution voxels as in DeepSTORM3D), resulting in over 20-fold improvement in prediction speeds, and the use of continuous sub-pixel coordinates eliminates a voxel size dependent limit on precision. The local output representation used by DECODE also avoids the potentially challenging non-local mapping of pixels to global coordinates used in DeepLoco.

Second, DECODE has four additional output channels that estimate the uncertainty of the localization along each coordinate given by *σxyz* = [*σx, σy, σz*] and of the brightness *σ N*. These predicted localization uncertainties can be used to filter out poorly localized detections to improve the rendering of super-resolution images. In addition, training the network to additionally predict the localization uncertainty corresponding to each detection also helps to improve the quality of the detection probabilities *p* by implicitly grouping all the detections corresponding to the same emitter. In contrast, standard output representations which only indicate the probability of detecting an emitter on a per-voxel basis make it more challenging to correctly group detection probability voxels corresponding to the same emitter in high emitter density and high localization uncertainty scenarios.

Third, the DECODE network integrates information across multiple frames with a two-stage design: The first stage (frame analysis module) analyses single imaging frames using a 2D multi-resolution convolutional network based on the “UNet” architecture^16^ to compute a feature representation of the single frame (1). The second stage (temporal context module) integrates the feature representations of the frame with those of the previous and next imaging frame using a second 2D UNet to produce the final predictions. As emitters persist over several frames, this improves detection and localization accuracy.

### Training the DECODE network using simulator learning

We train DECODE to simultaneously detect and localize emitters in SMLM measurements. Ground truth data for supervised learning are not easily available for SMLM. However, it is possible to simulate realistic images of activated emitters as the physics of imaging single molecules is well understood^8^. We train the DECODE network by generating a large amount of simulated data. To avoid structural bias^4^, we place emitters at random coordinates, and calculate simulated images with a realistic image-formation model that includes dye photo-physics, a measured PSF and camera noise (see Methods).

We trained the DECODE network to predict the probability of detection, along with the sub-pixel localization and localization uncertainty of each detected emitter. Our loss function has three terms: 1) a count loss that compares the true and detected number of emitters in the image; 2) a localization loss that trains the network to correctly localize the detected emitters and estimate the localization uncertainty and emitter brightness; 3) an optional background loss. The count and localization loss functions were derived together as an approximation to a spatial point process probability distribution. They work together to correctly train the DECODE network to predict one detection per emitter, and to correctly assign the localization uncertainty of each emitter to the corresponding detection. Together, they constitute a novel loss for counting, detecting, and localizing sets of discrete point-like objects.

The count loss first constructs a Gaussian approximation to the predicted number of emitters by summing the mean and the variance of the Bernoulli detection probability map, and then maximizes the probability of the true number of emitters under this distribution. Uncertain detections will lead to large predicted count variance, while confident detections will result in low variance. Thus, the count loss encourages a detection probability map with sparse but confident predictions. The localization loss models the distribution of sub-pixel localizations Δ*xyz* with a coordinate-wise independent Gaussian probability distribution^17^ with standard deviation *σxyz*. For imprecise localizations, this probability is maximized for large *σxyz*, for precise localizations for small *σxyz*. The distribution of all localizations over the entire image is approximated as a weighted average of individual localization distributions, where the weights correspond to the probability of detection. By optimizing both the probability of detection, the sub-pixel localization Δ*xyz*, and *σxyz* simultaneously, the network learns not only the best predictions for the coordinates of the emitters, but also the best estimate for their localization uncertainties. The emitter brightness predictions *N* and their uncertainties *σ N* are optimized similarly. Finally, the optional background loss computes the mean squared error between the true and predicted background images *B*.

### DECODE achieves high accuracy for a wide range of simulated data

#### Performance metrics

The quality of SMLM data analysis is commonly quantified by two factors: First, the detection accuracy quantifies the fraction of emitters that are detected. The metric we use here is the Jaccard Index (JI)^3^, that sets the true positives (TP) in relation to the false positives (FP) and false negatives (FN), JI = TP/(TP + FN + FP). The second factor is the localization error, i.e. how close the measured coordinates are to the true coordinates, measured here as the RMSE averaged over the dimensions (see Methods). We matched the detected emitters to the ground truth emitters in 3D with a lateral threshold of 250 nm and an axial threshold of 500 nm.

There is a natural trade-off between JI and localization error: Discarding all but the brightest and best separated emitters will result in a good (low) localization error but a bad (low) JI. Conversely, including also poorly localized emitters might improve JI, but deteriorates the localization error. The optimal operating point between these two extremes will depend on the experimental conditions and the scientific question. Because DECODE also provides uncertainties for each localization, it offers a straightforward way to filter localizations and thus set the desired balance between the number of detected emitters and the localization error that can be tolerated.

The Cramér-Rao Lower Bound (CRLB) gives the minimum achievable localization error for an optimal fitter given a known PSF, background, and noise model^18^. Most commonly, it is calculated under idealized conditions (i.e. non overlapping PSFs, homogeneous background, assuming the chosen PSF model to be the true model) and we use it here for comparison as a best-case limit for localization error.

#### DECODE approaches the Cramér-Rao Lower bound for low densities

We simulated 100, 000 frames with exactly one emitter per frame at random coordinates with a constant brightness and background and trained DECODE without temporal context. On this data with sparse activations, DECODE approaches the single emitter CRLB, i.e. the theoretical limit of precision (Fig. 2a). It thus performs as well as Maximum Likelihood Estimation (MLE) based fitters, which have also been shown to reach the CRLB^19^ in this regime.

**Figure 2.**
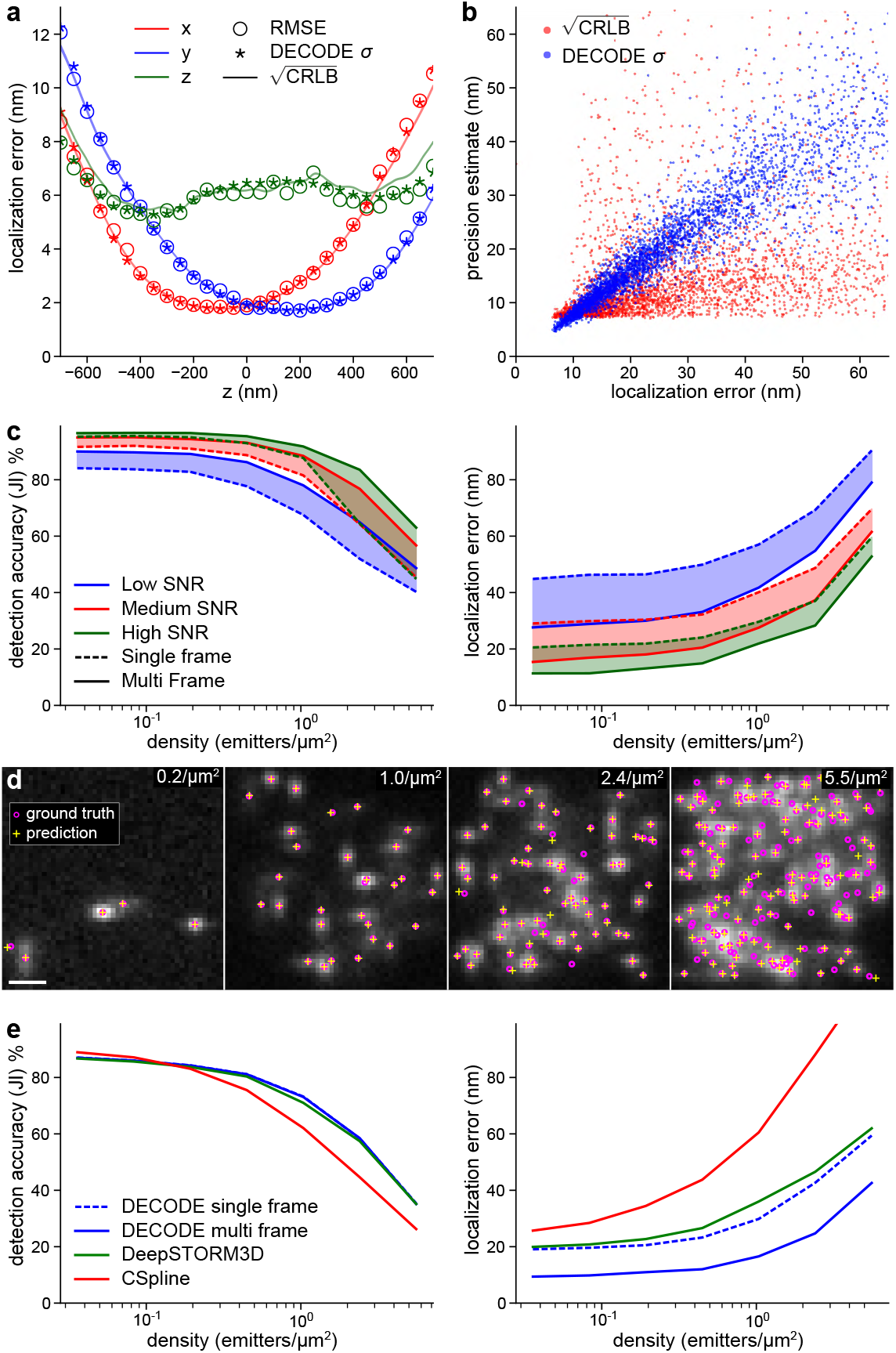
Performance of DECODE on simulated data: **a) DECODE reaches the single emitter Cramér-Rao Lower Bound (CRLB)** for isolated emitters. Root Mean Squarred Error (RMSE) and DECODE *σ* averaged over 50 nm bins. **b) The predicted localization uncertainty** *σ* **correlates well with the measured localization error for densely activated emitters.** We simulated the same dense emitter configuration 100 times and calculated the measured localization error as the RMSE of the predictions of the coordinates. In comparison, the (square-rooted) single-emitter-CRLB incorrectly under-estimates the true localization error for high emitter densities. **c) Temporal context improves both detection performance and localisation error.** We trained DECODE with (multi frame) and without temporal context (single frame) and compared detection accuracy and localization error for low, medium and high Signal to Noise Ratio (SNR). **d) Representative simulated frames** with ground truth coordinates (magenta circles) and predicted coordinates (yellow crosses) for the densities used in c and medium SNR. **e) Comparison of DECODE with CSpline and DeepSTORM 3D.** DECODE outperforms DeepSTORM and CSpline over a wide range of densities. See methods and Supplementary Table S1 for additional details on training and evaluation.

#### DECODE’s uncertainty estimates are well calibrated

In the high density regime, DECODE’s *σ* predictions correlate closely to the measured localization error (Fig. 2b), i.e. much better than the single emitter CRLB estimate which assumes isolated emitters (correlation coefficient 0.86 for *σ* vs. 0.07 for single emitter CRLB). For the low-density regime, the uncertainty estimates are in line with the measured error and the single emitter CRLB (Fig. 2a).

#### Temporal context improves localization error and detection

DECODE’s temporal context module pools information across multiple (we used 3) frames, to model the fact that emitters can persist in multiple subsequent frames. Use of this context module improves both the detection accuracy (JI) and the localization error (Fig. 2c). The increase in JI is apparent for all densities and SNRs. In addition, the RMSE is reduced by up to 20 nm. Overall, the temporal context has a large impact across imaging conditions, and is also more powerful than ‘grouping’ approaches which are often applied to localizations in a post-processing step (see Extended Data Fig. 2).

#### DECODE architecture outperforms a voxel based network architecture and a multi-emitter fitter

To assess how the DECODE network architecture performs against other deep learning based and iterative methods, we directly compared to DeepSTORM3D^13^ and CSpline^20^, a matching pursuit style multi-emitter fitter based on MLE, using the code provided by the authors. To minimise the risk of sub-optimal training, we trained DeepSTORM3D on data sampled from our generative model using the same parameters we used for the training of DECODE. For both DeepSTORM3D and CSpline we performed a parameter grid search over user-defined parameters to maximize their performance (measured as efficiency score^3^). To facilitate the comparison of localization precision, we filtered out DECODE localizations with the highest inferred uncertainties such that the remaining number match DeepSTORM3D. DECODE outperforms the other methods across all densities and SNRs (Fig. 2e, Extended Data Fig. 3) even without temporal context. When we use temporal context, DECODE reduces the localization error up to two-fold compared to DeepSTORM3D. Although both methods are based on deep learning, this performance improvement is based on the differences in output representation and loss function between DECODE and DeepSTORM3D. The localization error of DeepSTORM3D is limited by the super-resolution voxel size^13^ (Extended Data Fig. 4), which prevents the method from achieving the single emitter CRLB, unlike DECODE which has no such limitation. Notably, DECODE performs favourably in fitting time (Extended Data Fig. 5), taking less then 1.5 s to analyze 1000 frames of 64×64 pixels, while DeepSTORM3D requires between 34 s and 54 s and CSpline requires between 14 s and 2680 s, which is up to 1900-fold slower than DECODE.

#### DECODE outperforms all fitters on a public SMLM benchmark

The 2016 SMLM challenge is an on-going and continuously updated second generation comprehensive benchmark evaluation developed for the objective, quantitative evaluations of the plethora of available localization algorithms^3,21^. It offers synthetic datasets for training, created to emulate various experimental conditions. To avoid overfitting, evaluations are carried out on data not shared with contestants. It calculates various quality metrics, among them RMSE lateral or volume localization error, as applicable for 2D and 3D data respectively, the Jaccard index JI quantifying detection accuracy and a single ‘Efficiency’ score that combines RMSE and JI.

The performance of DECODE in the SMLM 2016 challenge, including extensive evaluations and side by side comparisons, is available online^†^. DECODE outperformed all 39 algorithms on 12 out of 12 datasets, often by a substantial margin (Fig. 3, data from challenge website, current as of Oct 1st, 2020). The datasets included high (N1) and low (N2, N3) signal to noise ratios (SNR), with low (LD) or high (HD) emitter densities, with 2D, astigmatism (AS) and double Helix (DH) PSF based imaging modalities.

**Figure 3.**
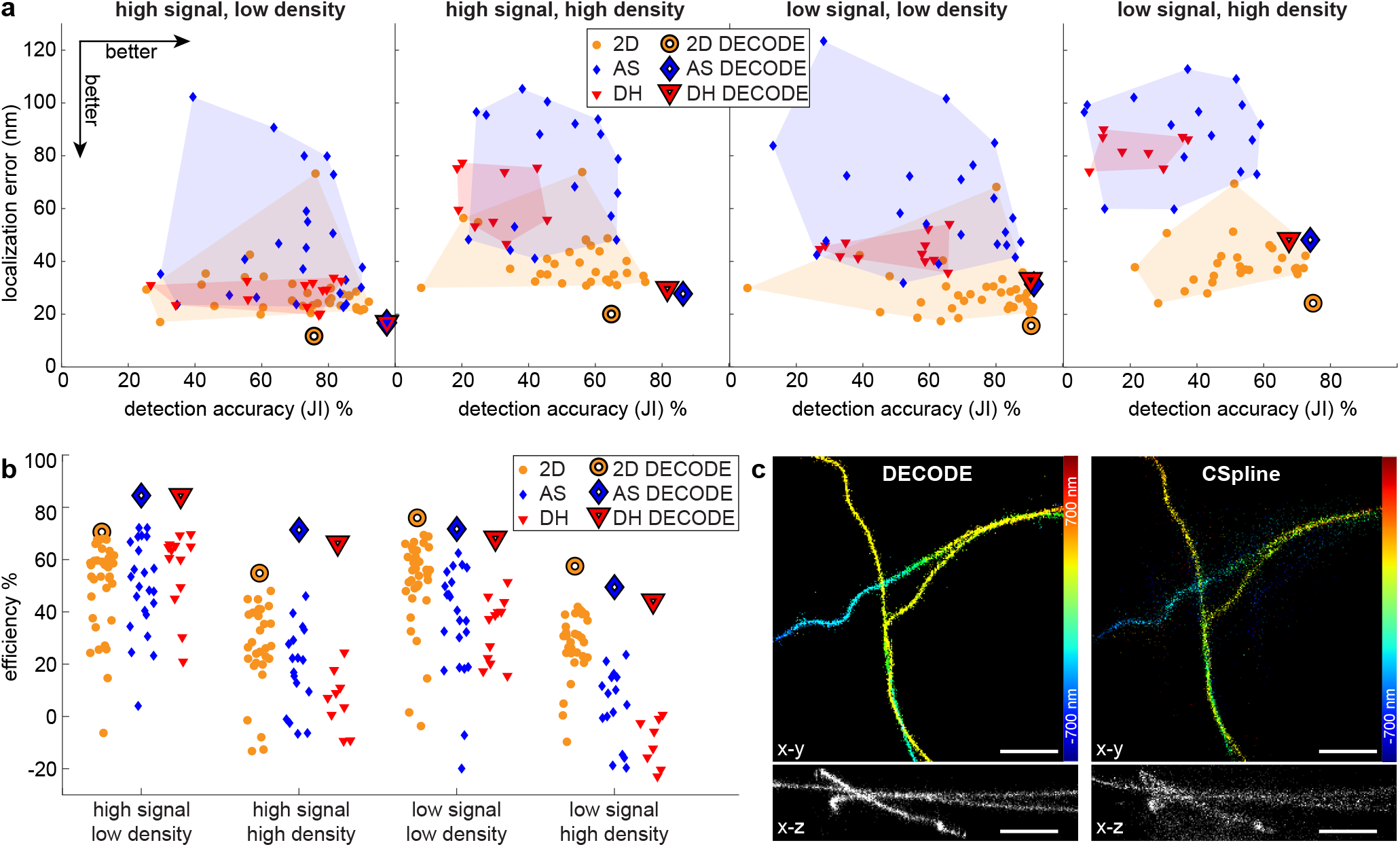
Performance comparison on the SMLM 2016 challenge. **a)** Performance evaluation on the twelve test datasets with low/high density, low/high SNR and different modalities (2D, AS: astigmatic, DH: double helix) using the detection accuracy (Jaccard Index, JI, higher is better) and localization error (lower is better) as metrics. Each marker indicates a benchmarked algorithm, large markers indicate DECODE. **b)** Efficiency scores (higher is better) for DECODE compared to other algorithms. Colored dots indicate performance numbers for other methods taken from the challenge website. **c)** Reconstructions by DECODE and the CSpline algorithm on the high density, low signal double helix challenge data. Upper panels *x*-*y* view, color coded by z coordinate, lower panels *x*-*z* reconstructions. Scale bars 1 μm.

DECODE achieves an average efficiency score of 66.6% out of the best possible score of 100% (achievable only by a hypothetical algorithm that accurately detects 100% emitters with 0 nm localization error). This is compared to an average score of 48.3% and 45.6% for all second and third place algorithms, respectively. The difference is particularly large under difficult imaging conditions, when high emitter densities and low SNR can conspire to make detection and localization challenging, particularly so for the double helix PSF. For example, compared to the second best algorithm (SMAP2018) in the Low SNR/high density/double helix condition, DECODE improves the localization error from 75.2 nm to 48.4 nm and the JI from 30.0% to 67.5%.

DECODE enhances super-resolution reconstructions by improving both the detection and the localization of single molecules. An example of this can be seen in Fig. 3c, where we compare the reconstruction obtained with DECODE CSpline^20^ on a high-density 3D double-helix dataset (using settings provided by the authors, github.com/ZhuangLab/storm-analysis). Thus, DECODE is setting new quantitative standards for localization algorithms, across both low and high SNRs and densities.

#### Considerations

As with any fitter, DECODE relies on an accurate PSF model and proper parameters, otherwise artifacts will dominate the predictions. When the localization uncertainty is large, for very dim and dense localizations far from the focal plane, DECODE has a bias towards predicting localizations close to the pixel center. This effect can be overcome by filtering out localizations with large predicted uncertainty (Extended Data Fig. 8, Methods).

### DECODE reduces imaging times by one order of magnitude

By enabling accurate emitter localization at high densities of more than 2.5 emitters per frame per μm^2^ (Fig. 2 c), DECODE can yield high-quality super-resolution reconstructions with much shorter imaging times. We demonstrate this by imaging and reconstructing the same sample of labeled microtubules at four different activation laser powers using STORM (stochastic optical reconstruction microscopy)^22,23^. This results in different emitter densities per frame. between 0.08 and 0.86 μm^−2^. The imaging time was chosen to result in the same number of total localizations and decreased from 1120 s to 460 s, 250 s and 93 s for stronger activation.

We trained and applied one common DECODE model to all four datasets (Fig. 4a). Whereas CSpline reconstructions quickly degrade with high emitter densities, DECODE consistently yields reconstructions with high accuracy even for the densest sample. We quantified the lateral resolution using Fourier Ring Correlation (FRC)^24^, which estimates resolution by measuring the correlation of two different reconstructions of the same image across spatial frequencies. DECODE consistently improves the x,y resolution by 20 nm - 30 nm over CSpline across all imaging densities (Fig. 4b and c) while detecting around 30% more localizations.

**Figure 4.**
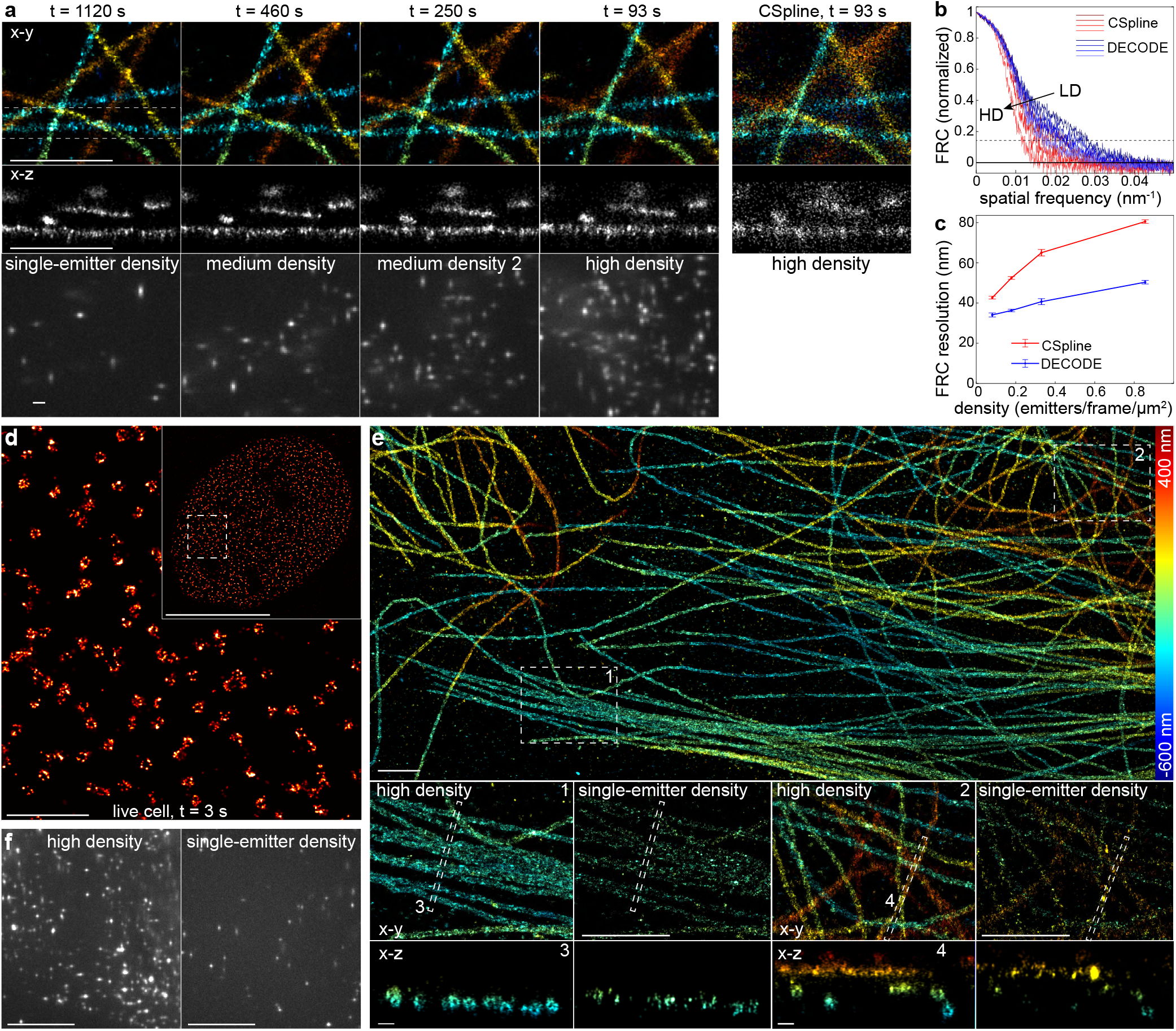
DECODE enables high-speed and live-cell SMLM and ultra-high labeling densities. **a) DECODE can reduce acquisition times by one order of magnitude.** The same sample of microtubules, labeled with anti-*α* tubulin primary and AF647 secondary antibodies, imaged with different UV activation intensities to result in different emitter densities between 0.08 and 0.86 emitters per frame per μm^2^ and acquisition times between 93 and 1120 s, while keeping the total number of localizations the same. For high-density activation, we show a comparison with CSpline. **b)** Fourier Ring Correlation curves for DECODE and CSpline for different emitter densities. **c)** Resolution estimates obtained using the Fourier Ring Correlation and 0.143 criterion across densities for both methods. **d) Fast live-cell SMLM** on the nuclear pore complex protein Nup96-mMaple acquired in 3 seconds. **e) DECODE enables ultra-high labeling densities.** Microtubules labeled with a high concentration of anti-*α* and anti-*β* tubulin primary and AF647 secondary antibodies. **e1, e2)** Magnified regions as indicated in a. Data acquired with high-density labeling shows continuous structures. As a comparison, the same sample was acquired after pre-bleaching of the fluorophores to reach the single-molecule blinking regime. Here, single labels are resolved in the superresolution reconstruction and lead to a sparse decoration of the microtubules. **e3, e4)** Side view reconstructions of regions as indicated in e1, e2 resolving the hollow, cylinder-like structure of immunolabeled microtubules. **f)** Representative raw camera frames for the high-density and single-emitter acquisitions, respectively. Scale bars: 10 μm (d inset, f), 1 μm (a, d, e, e1, e2), 100 nm (e3,e4).

### DECODE enables fast live-cell SMLM with reduced light exposure

Fast imaging is especially relevant for live-cell SMLM where the dynamics of the biological system under investigation dictate the necessary time resolution. At the same time, fast imaging usually requires high laser powers, deteriorates resolution^25^ and leads to substantial photo toxicity^26^. As DECODE allows activating emitters to high density, it enables faster imaging with decreased light dose for a given number of localizations. We were able to image nuclear pore complexes in living cells^27^ within only 3 seconds (Fig. 4d), 7 times faster than our previous speed-optimized live-cell SMLM^25^ and with 70% reduced light dose and thus photo toxicity.

### DECODE enables ultra-high labeling densities

Labeling densities in SMLM are fundamentally limited by the fraction of emitters that are in the bright state. For the best performing fluorophore Alexa Fluor 647, even without UV activation about 0.05% of the emitters are in the bright state^28^ due to activation by the red imaging laser and spontaneous activation. For the single-emitter blinking regime, this limits the number of emitters to about 200 per μm^2^. For higher labeling, pre-bleaching can be employed to reduce the number of emitters to this regime, but the resulting low labeling limits the resolution^15^ and in the superresolution reconstructions sparse individual emitters become dominant (Fig. 4e). With DECODE, we can now image densely labeled samples that previously were inaccessible. We demonstrated this on immuno-labeled microtubules that were labeled about 5-fold higher than compatible with single-emitter fitting, resulting in much smoother and denser decoration of the microtubules (Fig. 4e). In 50 nm thick orthogonal reconstructions, only the densely labeled microtubules were resolved as hollow cylinders, whereas after pre-bleaching to single-emitter blinking, these reconstructions only showed individual emitters (Fig. 4e,3,4).

### DECODE enables high fidelity reconstructions of 3D lattice light sheet PAINT

To illustrate the general applicability of DECODE, we applied it to 3D lattice light sheet (LLS) microscopy combined with the PAINT (point accumulation for imaging of nanoscale topography labeling) technique^15^. In PAINT microscopy, the fluorophore labeling a sample stochastically binds and unbinds from the sample, providing dense labeling. In LLS microscopy, thick volumes are imaged at high resolution by scanning a thin (1.1 μm) light sheet, with axial localization within the sheet enabled by astigmatism.

Single-molecule localization in LLS-PAINT is usually performed frame-wise using MLE fitting^29^. However, an emitter is visible in several adjacent z-planes in the volumetric data set. Thus, similar to exploiting the temporal context, we now use the same spatio-temporal context by analyzing three adjacent frames in the z-stack at the same time to improve detection accuracy and localization error.

We reconstructed a previously reported dataset of a chemically fixed COS-7 cell with intracellular membranes labeled by azepanyl-rhodamine (AzepRh)^15,29^ consisting of 70, 000 3D volumes comprising more than 10 million 2D images acquired in 270 nm steps. DECODE detected 500 million emitters, compared to 200 million emitters detected by the original algorithm. Thus, for a comparable quality of the reconstruction, only half of the frames are needed, reducing imaging times by over a day from 2.7 days to 1.35 days (Extended Data Fig. 6). At the same time, improved accuracy of DECODE results in sharper reconstructions (Fig. 5).

**Figure 5.**
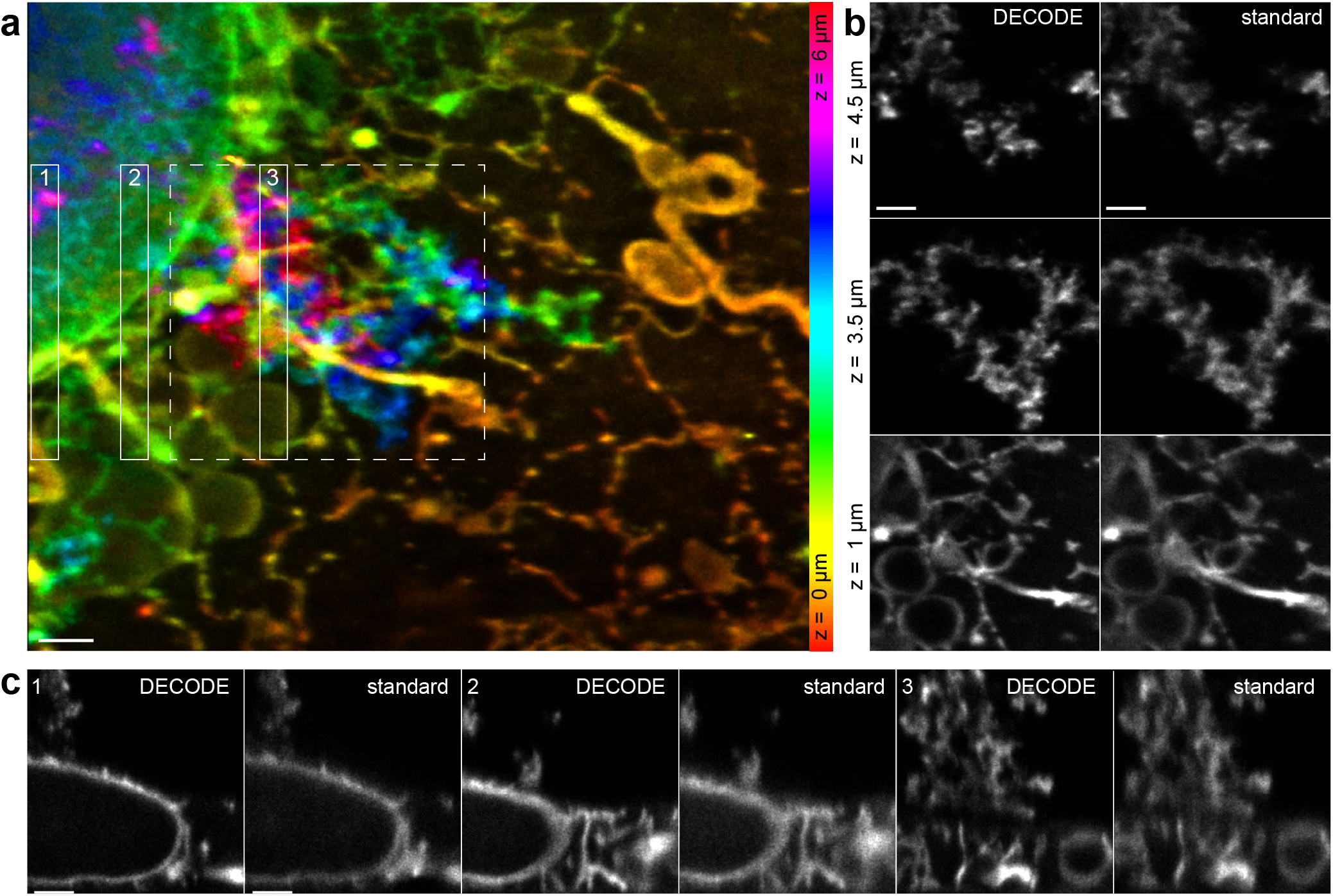
DECODE improves resolution in LLS-PAINT. **a)** COS-7 cell imaged with LLS-PAINT microscopy, overview. Data from Legant et al^29^, 70,000 volumes imaged over 2.7 days. **b)** 500 nm thick slices of the region indicated in a (dashed line), comparing DECODE analysis and the original analysis using MLE fitting (standard analysis). **c)** Perpendicular (side-view) reconstructions of 500 nm thick regions as indicated in a comparing DECODE and standard analysis. Scale bars 1 μm.

## Discussion

We presented DECODE, a new deep-learning based method for single molecule localization that performs exceptionally well on dense 3D data. DECODE differs from traditional localization algorithms by simultaneously performing detection and localization of emitters. It can be used in a flexible and general manner for a wide range of imaging parameters (including arbitrary Point Spread Functions and noise models) and imaging modalities such as 3D lattice light sheet PAINT imaging. In a publicly available benchmark challenge it is the best performing algorithm in every condition, and often improves both localization and detection accuracy by a large margin.

By making use of the temporal context, DECODE improves detection accuracy and localization error of emitters that are active across multiple imaging frames. Temporal context is also used by post-processing steps in SMLM relying on ‘merging’ or ‘grouping’ of localizations, in which localizations occurring in consecutive images that are closer to each other than a fixed threshold are assumed to belong to the same emitter and their coordinates are averaged, weighted by the uncertainty of each localization. However, grouping does not improve detection of emitters, and it fails for dense or dim emitters whose localizations cannot be linked unambiguously across frames.

DECODE not only predicts coordinates of emitters, but also their uncertainty. This is highly useful for filtering out imprecise localizations, for reconstruction of superresolution images in which every localizations rendered as a Gaussian with a size proportional to the coordinate uncertainty, and as weights for quantitative coordinate-based analysis of SMLM data.

We demonstrated the performance of DECODE on various experimental SMLM data sets. We could show that the excellent performance on high-density data can increase the achievable localization density or decrease imaging times by one order of magnitude. This allowed us to perform live-cell measurements on nuclear pore complexes with high temporal resolution and reduced light exposure, and to achieve ultra-high labeling on microtubules. LLS-PAINT data analyzed with DECODE showed markedly improved resolution due to substantial improvements in emitter detection and localization error.

Prediction of coordinates with DECODE can be as fast as GPU-based MLE-fitters for sparse activation, but greatly outperforms those for high densities, as the computational complexity of DECODE depends only on the size of the image and not the number of emitters in each imaging frame. However, it requires the training of a new neural network whenever the optical properties of the microscope change. This training can currently take over 10 hours on a single GPU, but after just 2 hours of training time, the localization error is within 1 nm and the JI within 2% of the final value (Extended Data Fig. 7). To reduce training times further, one can likely take an existing network and fine-tune its parameters using a smaller number of simulations, rather than training it from scratch. Ultimately, it may be possible to train a single network across multiple parameter settings or even PSFs, so that the same network can ‘amortize’ inference across multiple experimental settings.

To make DECODE easily usable by the entire community, we distribute it as a Python-based open source software package based on the PyTorch^30^ deep learning library. We provide pre-compiled, easily installable code, along with detailed tutorials and integration into the SMAP SMLM analysis software^31^. To enable anyone to directly use DECODE for training and prediction without relying on prior programming knowledge and dedicated local hardware, we deploy these Jupyter notebooks in Google Colab, complementing a recent initiative to make deep learning based image analysis tools accessible to non-experts at minimal cost^32^. Thus, DECODE will enable a large community to directly perform SMLM in a new high-density regime with greatly increased imaging speeds or localization densities and excellent localization and detection accuracy.

## Data availability

All data can be publicly accessed from the DECODE webpage (see Code Availability).

## Code Availability

DECODE is available at http://github.com/TuragaLab/DECODE.

## Acknowledgments

S.C.T. is supported by the Howard Hughes Medical Institute, J.R. and L.-R.M were supported by the European Molecular Biology Laboratory, the European Research Council (CoG-724489 to J.R.) and the the National Institutes of Health Common Fund 4D Nucleome Program (grant no. U01 EB021223 to J.R.). J.H.M. and A.S. were supported by the German Research Foundation (DFG) through Germany’s Excellence Strategy (EXC-Number 2064/1, Project number 390727645) and the German Federal Ministry of Education and Research (BMBF, project ‘ADMIMEM’, FKZ 01IS18052). We thank Daniel Sage for useful discussions, Ulrike Boehm, David Greenberg and Poornima Ramesh for comments on the manuscript, and Eric Betzig and Jennifer Lippincott Schwartz for kindly sharing data with us.

## Author contributions

A.S., L.-R.M., J.H.M., J.R. and S.C.T. conceived the project, analyzed the results and wrote the manuscript with input from all authors. A.S. and L.-R.M. wrote the software, U.M. and J.R. acquired the biological data, A.K provided supervision, C.J.O. and W.R.L. provided the LLS data and helped with analysis.

## Methods

### DECODE network architecture for probabilistic single molecule detection and localization

Our architecture consists of two stacked U-nets^16^ (Extended Data Fig. 1), each with two up- and downsampling stages and 48 filters in the first stage. Each stage consists of three fully convolutional layers with 3 × 3 filters. In each downsampling stage, the resolution is halved, and the number of filters is doubled, vice versa in each upsampling stage. Upsampling is performed using nearest neighbor interpolation to avoid checkerboard artifacts^33^. For multi-frame DECODE, three consecutive frames are processed by the the first frame analysis U-net (with parameters shared for every frame), and the outputs are concatenated and passed to the second temporal context U-net. The entire DECODE network is always trained end-to-end by gradient descent.

For each camera pixel *k*, the DECODE network predicts i) a Bernoulli probability map *p*_*k*_ that an emitter was detected near that pixel, ii) the coordinates of the detected emitter Δ*x*_*k*_, Δ*y*_*k*_, Δ*z*_*k*_ relative to the center of the pixel *x*_*k*_, *y*_*k*_*z*_*k*_ , iii) a non-negative emitter brightness (“photon count”) *N*_*k*_, and iv) the uncertainties associated with each of these predictions, *σx*_*k*_, *σy*_*k*_, *σz*_*k*_, *σ N*_*k*_. For each of these outputs, we use two additional convolutional layers that follow the second U-net. We used the Exponential Linear Unit (ELU) activation function^34^ for all hidden units, and the logistic sigmoid nonlinearity for the non-negative detection probability *p*, brightness *N*, and the uncertainty outputs *σx, σy, σz, σ N* (scaled by a pre-factor of three). For the coordinate outputs Δ*x*, Δ*y*, Δ*z* we use the hyperbolic tangent nonlinearity which limits their range to [−1, 1] (i.e. to twice the size of a pixel). This way, even though the network can at most predict one emitter per pixel, when necessary the neighboring pixels can each contribute in order to place multiple localizations within a single pixel.

### A loss function for simultaneous detection, localization, and uncertainty estimation

Given a set of *E* simulated emitters active in each imaging frame with locations for each emitter *e* given by *x*_*e*_, *y*_*e*_, *z*_*e*_ and brightness *N*_*e*_, and a background image map *B*_*k*_ simulated as described below, we developed a loss function that trains the DECODE network to detect the correct number of emitters, to predict the sub-pixel localization and brightness for each detection (along with the uncertainty), and to predict the image background. Our loss function is a sum of three terms – a count loss 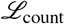, a localization loss 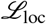, and a background loss 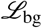,

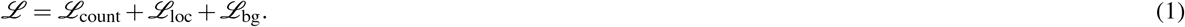

The **count loss** 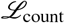 is a function of the detection probability map *p*_*k*_ with *K* total pixels and the total number of true emitters *E*. Interpreting *p*_*k*_ as a Bernoulli detection probability for a single emitter, we can compute the mean and variance of the predicted total number of emitters detected, if we were to independently sample binary detections from each *p*_*k*_. While the predicted count distribution *P*(*E*|{*p*_*k*_}) over the number of emitters detected by this Bernoulli sampling procedure follows an intractable Poisson Binomial distribution, we can approximate this predicted distribution as a Gaussian distribution,

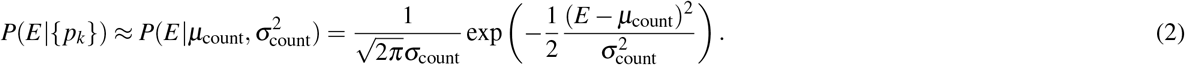

The mean of a sum of Bernoulli random variables is the sum of the means 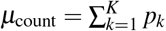, and the variance is the sum of the variances of each independent Bernoulli random variable 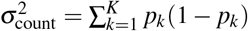. This count loss maximizes the log probability of the true number of emitters *E* under the Gaussian approximation of the predicted count probability distribution. This loss is minimized when *μ*_count_ correctly matches *E*, sparsely predicting only one non-zero *p*_*k*_ per detected emitter, and when 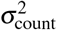 is small, which happens when *p*_*k*_ are confident and so nearly binary,

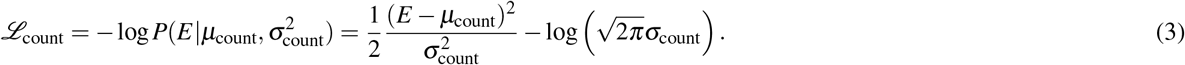

The **localization loss** 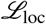 is a function of the the true emitter locations, and the predicted detection probability map, and the sub-pixel localizations Δ*x*_*k*_, Δ*y*_*k*_, Δ*z*_*k*_, brightness *N*_*k*_, along with the associated uncertainties *σx*_*k*_, *σy*_*k*_, *σz*_*k*_, *σ N*_*k*_ for each detected emitter. For each pixel *k*, we predict a 4D Gaussian distribution 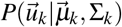 over the the absolute position and brightness of an emitter 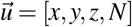 detected in pixel *k* corresponding to the mean and uncertainty in the sub-pixel localization and brightness of the emitter detected in pixel *k*, with mean 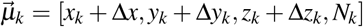, and diagonal covariance matrix 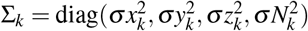,

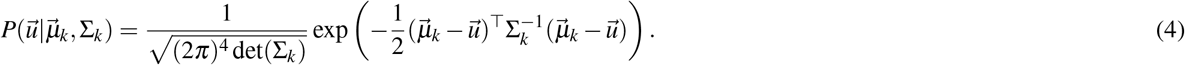

Here, the *x*_*k*_, *y*_*k*_, and *z*_*k*_ are the absolute coordinates for the center of pixel *k*, so *x*_*k*_ + Δ*x*_*k*_ corresponds to the absolute coordinates of the emitter to sub-pixel precision. We note that the localization loss defined below ignores the predicted localization and brightness for pixels where no emitter is detected, i.e. *p*_*k*_ is zero.

At any given point in training, the true number of emitters will not necessarily match the detected number of emitters perfectly, and we will not have a perfect correspondence between predicted emitters and true emitters. A full probabilistic loss function would sum over all possible assignments of true emitters to detected emitters in order to correctly evaluate 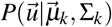. And since *p*_*k*_ will not necessarily be sparse, the correct cost function would include an intractably large sum over 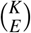 terms. We approximate this by constructing a Gaussian mixture model over the predicted per pixel distributions 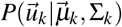 with mixture weights equal to 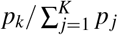 where the denominator is a sum of the detection probability over all pixels in the image.

The resulting approximation leads to the following localization loss function which maximizes the probability of the true absolute coordinates and brightness of each ground truth emitter 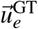 under the weighted mixture of per pixel probabilities,

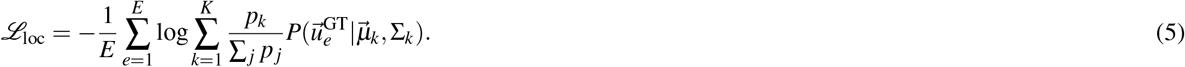

The **background loss** 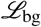 computes the simple squared error between the predicted and true background maps,

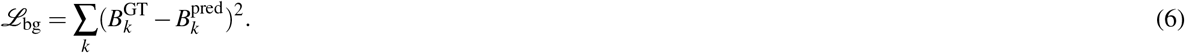

### Obtaining localizations and post-processing

The DECODE network predicts the probabilities *p*_*k*_ of an emitter being located at a specific pixel *k*. To get deterministic, fast and precise final localizations we use a variant of non-maximum suppression^35^. First, to get a binary mask of emitter candidates for a given frame we identify probability peaks, i.e. pixels with values that are above 0.3 and higher than all values in a surrounding 3 × 3 patch. We then add the probability mass from the 4 directly adjacent pixels to the values at the candidate coordinates by convolving the probability map with a cross shaped filter and applying the mask. All candidates with added probability values above 0.7 are counted towards the localizations. The algorithm can be expressed purely in the form of pooling and convolution operations and therefore runs efficiently on a GPU.

For difficult imaging conditions when the predicted localization uncertainties are large, i.e. high densities, low SNR values, and large offsets from the focal plane, the sub-pixel coordinates Δ*x*, Δ*y*, and Δ*z* can be biased towards the center of the pixels (Extended Data Fig. 8). This is because with large predicted localization uncertainty, the predicted mean location is poorly constrained. This bias towards 0 (pixel center) scales with the uncertainty of the predictions and can produce artifacts in the reconstructed image depending on how the reconstruction is performed. If a reconstruction uses only the coordinates while ignoring the uncertainty, poorly localized emitters will cluster towards the pixel centers. A more expensive rendering procedure which renders a Gaussian localization distribution with variance proportional to the estimated uncertainty corresponding to each emitter will reduce the impact of this artifact since the bias is usually small relative to the localization uncertainty. Also, filtering out localizations with high uncertainty removes this artifact (Extended Data Fig. 8).

### Simulating training data

Training samples are continuously generated in an asynchronous fashion and each frame is only used once as a target. For this reason the network cannot overfit to specific frames. The performance of our approach will depend on an accurate generative model and could show reduced performance when there is a mismatch between the simulated and experimental data. Thus, we developed a realistic model for the image formation process that incorporates dye blinking behaviour, a realistic PSF model and realistic camera read noise.

#### Structural prior

While incorporating prior structural information has shown to be beneficial^36,37^, there are concerns that these priors could potentially bias the model to the training data, which could result in the presence of misleading structures after the fitting procedure. We therefore sample the coordinates of the emitters from a 3D homogeneous spatial Poisson point process distribution with density as specified in the text, limits corresponding to the size of the image and the z-range for which the PSF was calibrated.

#### Photophysical prior

In contrast to prior work, DECODE can directly incorporate temporal context into the detection and localization of emitters, rather than as a post-processing step. We simulate the temporal dynamics of emitters, at least over the short time scale of three imaging frames corresponding to the temporal context of the DECODE network.

For each emitter, the time of initial appearance *t*_0_ is sampled from a continuous random distribution. The on-time of the emitter follows an exponential distribution parametrized by *λ*. For each emitter, we draw a photon flux from a Gaussian distribution *N*(*μ*_flux_, *σ*_flux_). Together with the amount of time the emitter is active in each frame this determines the total number of photons emitted in a frame. Since the input to our model is only a window of three frames, we argue that it is not necessary to model long range temporal correlations that are part of a more detailed photo activation model^38^, like an emitter in the dark state which reappears many frames later. The aforementioned parameters are estimated by a *prefit* procedure as described in *Estimating simulation parameters*.

#### Point spread function

The PSF is a fundamental characteristic of a microscope, specifying the image formed by a single point emitter, and we approximate it to be spatially invariant across the field of view. Given the object 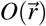 in the object plane, and 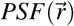, the image 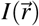 results as

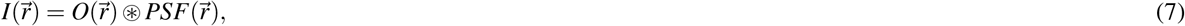

where ⊛ denotes the convolution operator. While Gaussian approximations of the PSF are frequently used for both 2D and 3D^39,40^ data, (cubic) spline functions have been shown to achieve more accurate results and can mimic almost arbitrary PSF’s^19,20^. Following *Li et al.*^19^ and *Babcock et al.*^20^ a three-dimensional PSF can be modelled as

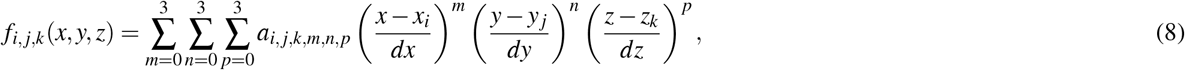

where *i, j, k* are the voxel indices, *dx, dy* are the pixel sizes; *dz* is the step size in the axial dimension; *x*_*i*_, *y*_j_, *z*_*k*_ are the corner coordinates of the voxel (*i, j, k*) in the respective directions and *a*_*i,j,k,m,n,p*_ are the respective spline coefficients, which amounts to 64 coefficients per pixel and per z-slice. In a bead calibration routine, the coefficients *a*_*i,j,k,m,n,p*_ are estimated and account for varying experimental conditions. Because of the simple form of equation 8, the CRLB with respect to the fitting parameters *x, y, z* can be calculated easily as the diagonal elements of the inverse of the Fisher information matrix.

#### Camera model

All real datasets presented in this work were recorded with an EMCCD camera, with the exception of the LLS data which was recorded with an sCMOS camera. The measured camera signal is subject to various noise sources, which we will discuss in the following:

***Shot noise*** originates from the stochastic nature of photons when interacting with the camera chip. The expected number of detected electrons is

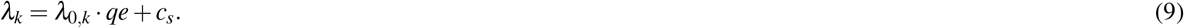

Here, *λ*_0,*k*_ is the expected number of photons that are collected in pixel *k*, *qe* is the quantum efficiency, and *c*_*s*_ the spurious charge, measured in electrons. The probability *p*_shot_(*s*_*k*_) of observing the signal *s*_*k*_ in pixel *k* follows a Poisson distribution,

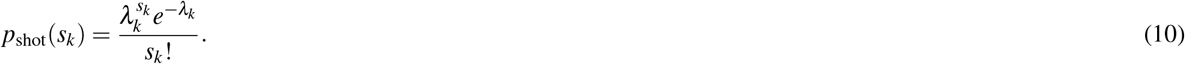

***EMCCD Amplification noise*** stems from the amplification of photo electrons that pass through the gain register and stochastically generate additional electrons. For our EMCCD camera noise model we follow *Huang et al.*^41^. EMCCD amplification noise can be described approximately by a Gamma distribution,

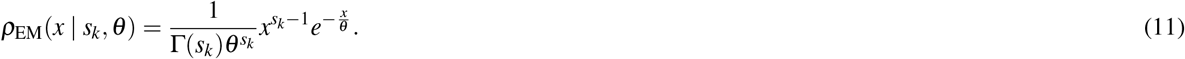

*ρ*_EM_(*x*|*s*_*k*_, ***θ***) denotes the probability that *s*_*k*_ input photo electrons in pixel *k* with an EM gain of ***θ*** create *x* output electrons after the gain register.

***Read noise*** stems from the process of converting electrons into a digital signal. In this process, the signal is usually multiplied by a gain factor *g* and an offset *o* is added to avoid negative signal. In this work, we convert the input camera image to photon units prior to inference by subtracting *o* and dividing by *g*. In addition, when using EMCCD cameras we divide by the EM gain ***θ***, thus the units of the read noise are photo electrons. We approximate the read noise (both for sCMOS and EMCCD cameras) by a zero mean additive Gaussian distribution with variance *σ*^2^,

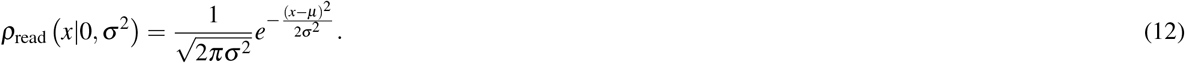

### Training details

Training was performed on 40×40 pixel sized regions that are directly simulated or randomly selected from larger simulated images at each iteration. We used the AdamW optimizer^42^ with a group learning rate of 6 × 10^−4^ for the network parameters. We reduce the learning rate by a factor of 0.5 if the relative change of the validation loss is less than 10^−4^ for at least 10,240 iterations with a batch size of 64. To stabilize training we employ gradient norm clipping with a maximum norm of 0.03. Very dim emitters with less then 50 photons are excluded from the ground truth targets (but still rendered) so that the network is discouraged to make predictions for practically invisible emitters.

#### Estimating simulation parameters

For training DECODE, a proper parametrization of the simulation is needed to match the real data distribution. In a prefitting step, the main parameters, i.e. the emitter on-time, emitter brightness and background, can be determined. The prefitting can be performed with a single-emitter MLE fitter after filtering the log-likelihood value to exclude data from overlapping PSFs. This step is incorporated in the SMAP software for the sake of ease of use^31^. We observed that the precise values of the simulation parameters of the emitters’ photophysics (i.e. lifetime and brightness) and density are not crucial, as the stochastic nature of the emitters’ position, brightness and appearance time presents the network with data that matches the real experiments under different conditions and effectively covers a broad range of these parameters. The camera parameters are usually given by the manufacturer. The given network architecture and training parameters are effective across different real and simulated datasets and in our experience do not have to be optimized by the end user.

### Evaluating localization error and reconstruction resolution

To evaluate performance on the challenge datasets, as well as our own simulations, we use two metrics:

First, the **localization error**, measured in nm, is the RMSE averaged over the dimensions:

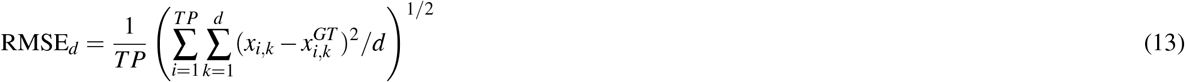

*TP* is the number of localizations that are matched to ground truth coordinates, *d* is the dimension (2 for 2D data, 3 for 3D data), *x*_*k*_ = *x, y, z* are the predicted coordinates and 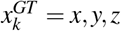 the ground truth coordinates.

Second, the **detection accuracy** or Jaccard index JI, which quantifies how well an algorithm does at detecting all the emitters while avoiding false positives:

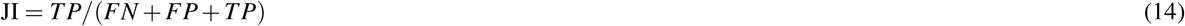

*T P* are the true positives, *FN* the false negatives and *FP* the false positives.

Localizations are matched to ground truth coordinates when they are withing a circle of 250 nm radius and the distance in *z* is less than 500 nm. As a single metric that evaluates the ability to reliably infer emitters with high precision we use the efficiency metric as defined in Ref. 3:

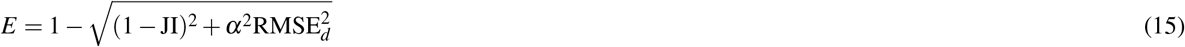

Lateral and axial efficiency are calculated with alpha values of *α* = 1 × 10^−2^ nm^−1^ and *α* = 0.5 × 10^−2^ nm^−1^ respectively and then averaged to obtain the overall efficiency. Detection accuracy is expressed in units of 0 to 1 (or 0% to 100%), the efficiency ranges up to 1 (or 100%)· for a perfect fitting algorithm.

The Fourier ring correlation^24,43^ (FRC) in Fig. 4 was calculated by constructing two super-resolution images of the same sample (pixel size 10 nm) by dividing the localizations into two sets. We did this by alternating blocks of 50k consecutive localizations.

### Simulating data for performance evaluation

To simulate data for performance evaluation and comparison shown if Fig. 2 we assumed an ideal camera without EMCCD or read noise and an image size of 64 × 64 pixels. We used the PSF model that was acquired for the data set in Fig. 4a. Data used to test the effect of the SNR and density were simulated using the structural and photophysical prior previously described with an average on-time of 2 frames. Precise simulation parameters can be found in Supplementary Table S1. The CRLB is evaluated as the diagonal elements of the inverse of the Fisher information matrix^18^ with the fitted parameters and spline interpolated experimental PSF model and was calculated with the SMAP software^31^.

### Comparison with DeepSTORM3D and CSpline

For both methods we used the software provided by the authors. For the DeepSTORM3D comparison instead of using their PSF fitting procedure and generative model we sampled ground truth coordinates and training images using our model so that it exactly matches the simulated test data. To minimize possible effects of overfitting we generated 22,500 images with a size of 121 × 121 pixels (22k for training and 500 for validation). DeepSTORM3D uses a fourfold super-resolved grid in the *x − y* dimensions and we chose discretization of 15 nm in *z*. Therefore each voxel of the output represenation has a size of 30 × 30 × 15 nm. For DECODE (with and without temporal context), and DeepSTORM3D we trained six networks each on training data generated with average emitter densities of 0.65 and 2.17 μm^−2^ as well as low, medium and high SNRs (1000, 5000 and 20,000 average photons). We used the low density network for the CRLB evaluation (Fig. 2a, Extended Data Fig. 4) and the simulated data with densities between 0.04 and 2.4 μm^−2^ (Fig 2c,d, Extended Data Fig. 3) and the high density networks for densities between 2.4 and 5.6 μm^−2^. DeepSTORM3D has two hyperparameters that control the post-processing and determine the balance between recall and localization error. We performed a sweep over combinations of radius = [5,6,7,8,10] and threshold = [5,8,12,20,30,40] and picked the values that maximized the efficiency score on the validation data for each of the six networks. We discovered and fixed a bug in the DeepSTORM3D post-processing software which lead to poor localizations. All DeepSTORM3D results were reported with the fixed post-processing algorithm.

For the CSpline comparison we created a bright artificial bead with 500k photons using our PSF model, which we used to generate the CSpline PSF model. The most critical settings are the find-max-radius and threshold, which we again optimized by sweeping over values find-max-radius = [[2,3,4,5], threshold = [6,7,8,9,10] to maximize efficiency for each of the three SNRs on data generated with an average emitter densities of 0.9 μm^−2^.

### DECODE for LLS-PAINT microscopy

A DECODE model for lattice light sheet point accumulation for imaging of nanoscale topography (LLS-PAINT) microscopy^15^ was trained by simulating the imaging of an angled light sheet being swept through a volume. This leads to the same emitter appearing with fixed shift in the *x* and *z* coordinates relative to the imaged plane between consecutive camera frames. The offset in emitter coordinates from frame to frame are given by the microscope geometry as described in^29^. We simulated data with high emitter densities to match the densities seen in LLS-PAINT.

We analyzed a large dataset corresponding to a fixed COS-7 cell with intracellular membranes labeled with azepanyl-rhodamine (AzepRh) described in Legant et al.^29^. Over a period of 2.7 days (64.8 hours), LLS-PAINT imaging yielded 70,000 3D volumes comprising more than 10 million 2D images. Significant non-uniform swelling of the sample was observed over the course of the imaging, which was approximately corrected by non-rigid registration in Legant et al.^29^. We applied the same correction transformation estimated by Legant et al.^29^ to DECODE localizations.

We introduced an additional simulation-free training step and loss function to the training of the LLS-PAINT DECODE network based on the Re-weighted Wake Sleep algorithm^44^ for training variational autoencoders (VAE)^45,46^. This form of auto-encoder learning allowed us to further optimize the parameters of the PSF and improve the background predictions based on the real data, as opposed to the simulation.

### Sample preparation

#### Sample seeding

Before seeding of cells, high-precision 24 mm round glass coverslips (No. 1.5H, catalog no. 117640, Marienfeld) were cleaned by placing them overnight in a methanol:hydrochloric acid (50:50) mixture while stirring. After that, the coverslips were repeatedly rinsed with water until they reached a neutral pH. They were then placed overnight into a laminar flow cell culture hood to dry them before finally irradiating the coverslips by ultraviolet light for 30 min. Cells were seeded on clean glass coverslips 2 days before fixation to reach a confluency of about 50 – 70% on the day of fixation. They were grown in growth medium (DMEM (catalog no. 11880-02, Gibco)) containing 1× MEM NEAA (catalog no. 11140-035, Gibco), 1× GlutaMAX (catalog no. 35050-038, Gibco) and 10% (v/v) fetal bovine serum (catalog no. 10270-106, Gibco) for approximately 2 days at 37 °C and 5% CO2.

#### Sample staining

##### Preparation of Nup96-mMaple samples

For imaging of live cells, coverslips containing Nup96-mMaple cells^27^ (catalog no. 300461, CLS Cell Line Service, Eppelheim, Germany) were rinsed twice with warm PBS before they were mounted onto a custom manufactured sample holder in 1 mL growth medium containing 20 mM HEPES buffer and imaged directly.

##### Preparation of microtubule samples

For microtubule staining, wild-type U-2 OS cells (ATCC HTB-96) were prefixed for 2 min with 0.3% (v/v) glutaraldehyde in cytoskeleton buffer (CB, 10 mM MES pH 6.1, 150 mM NaCl, 5 mM EGTA, 5 mM glucose, 5 mM MgCl2) + 0.25% (v/v) Triton X-100 and fixed with 2% (v/v) glutaraldehyde in CB for 10 min. Fluorescent background was reduced by incubation with 0.1% (w/v) NaBH4 in PBS for 7 min. After samples had been washed three times with PBS, microtubules were stained with anti-*α*-tubulin (MS581; NeoMarkers, Fremont, CA, USA), and for ultra-high labeling (Fig. 4e) additionally with anti-*β* -tubulin (T5293; Sigma-Aldrich), each diluted 1:50 in PBS with 2% (w/v) BSA, overnight. After being washed three times with PBS, samples were incubated with anti-mouse Alexa Fluor 647 (A21236; Invitrogen, Carlsbad, CA, USA) 1:50 in PBS + 2% (w/v) BSA for 6 h. After being washed three times with PBS, samples were imaged in blinking buffer as described above. The holder was sealed with parafilm.

### Microscopy

SMLM data were acquired on a custom built widefield setup described previously^47,48^. Briefly, the free output of a commercial laser box (LightHub, Omicron-Laserage Laserprodukte) equipped with Luxx 405, 488 and 638 and Cobolt 561 lasers and an additional 640 nm booster laser (iBeam Smart, Toptica) were coupled into a square multi-mode fiber (catalog no. M103L05). The fiber was agitated as described in Ref. 49. The output of the fiber was magnified by an achromatic lens and focused into the sample to homogeneously illuminate an area of about 400 μm^2^. The laser is guided through a laser cleanup filter (390/482/563/640 HC Quad, AHF) to remove fluorescence generated by the fiber. The emitted fluorescence was collected through a high numerical aperture (NA) oil immersion objective (HCX PL APO 160×/1.43 NA, Leica), filtered with a 676/37 (catalog no. FF01-676/37-25, Semrock) bandpass filter (for imaging of Alexa Fluor 647) or with a 600/60 (catalog no. NC458462, Chroma) bandpass filter (for live-cell imaging of Nup96-mMaple) on an EMCCD camera (Evolve 512, Photometrics). Astigmatism was introduced by a cylindrical lens (f = 1.00 m; catalog no. LJ1516L1-A, Thorlabs) to determine the z coordinates of fluorophores. The z focus was stabilized by an infrared laser that was totally internally reflected off the coverslip onto a quadrant photodiode, which was coupled into closed-loop feedback with the piezo objective positioner (Physik Instrumente). Laser control, focus stabilization and movement of filters was performed using a field-programmable gate array (Mojo, Embedded Micro). The pulse length of the 405 nm laser can be controlled by a feedback algorithm to sustain a predefined number of localizations per frame.

Coverslips containing prepared samples were placed into a custom build sample holder and 500 μL of blinking buffer (50 mM Tris/HCl pH 8, 10 mM NaCl, 10% (w/v) d-glucose, 500 μg/mL glucose oxidase, 40 μg/mL glucose catalase, 35 mM MEA) was added for imaging of Alexa Fluor 647 samples. Live-cell imaging of Nup96-mMaple was performed in PBS in D2O containing 10% 10x PBS in 90% (v/v) D2O.

For live-cell imaging of Nup96-mMaple (Fig. 4d), we used an excitation intensity at 561 nm of 16.7 kW/cm^2^ and a UV laser power of 80 W/cm^2^. The exposure time was 12 ms and the pulse length of the UV laser was automatically adjusted from 1 ms to 12 ms to keep the density of localizations constant.

For imaging of microtubules at different activation densities (Fig. 4), we used an exposure time of 15 ms and an excitation intensity at 640 nm of 15.5 kW/cm^2^. We adjusted the UV pulse length to result in the desired density of activated fluorophores. As we started with the highest density, by the time we imaged the lowest density a large fraction of the fluorophores was bleached so that we could operate in the single-emitter regime.

For imaging microtubules with ultra-high labeling, we used an exposure time of 15 ms and an excitation intensity at 640 nm of 13.4 kW/cm^2^ and no UV activation.

## Extended Data Figures

**Extended Data Figure 1.**
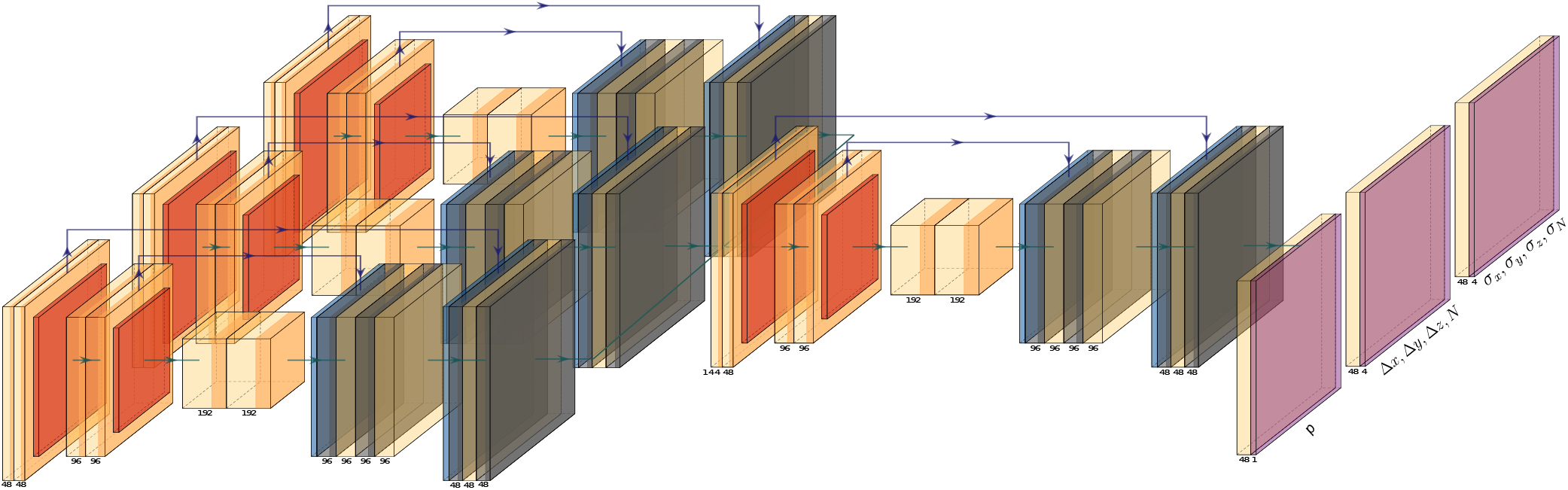
Architecture. The DECODE network consists of two stacked U-Nets^16^ with identical layouts (the three networks depicted on the left share parameters). The *frame analysis module* extracts informative features from three consecutive frames. These features are integrated by the *temporal context module*. Both U-Nets have two up- and downsampling stages and 48 filters in the first stage. Each stage consists of three fully convolutional layers with 3 × 3 filters. In each downsampling stage, the resolution is halved, and the number of filters is doubled, vice versa in each upsampling stage. Blue arrows show skip connections. Following the *temporal context module* three output heads with two convolutional layers each produce the output maps which have the same spatial dimensions as the input frames. The first head predicts the Bernoulli probability map *p*, the second head the spatial coordinates of the detected emitter Δ*x*, Δ*y*, Δ*z* and its intensity *N* and the third head the associated uncertainties *σx, σy, σz, σ N*. An optional fourth output head can be used for background prediction.

**Extended Data Figure 2.**
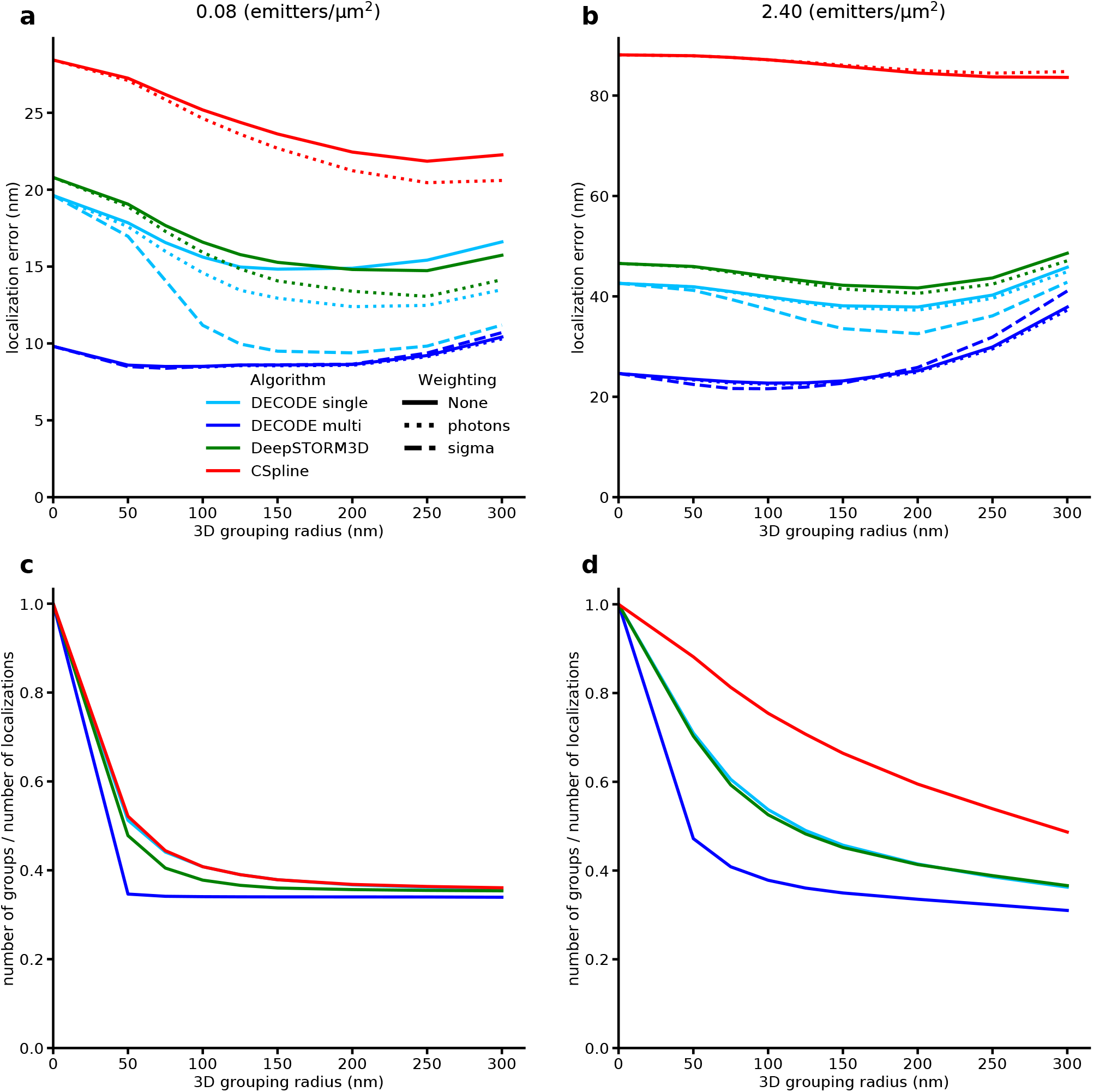
Impact of grouping across grouping radius for different averaging weights. Predictions in consecutive frames are grouped when they are closer to each other than the given grouping radius. A grouping radius of 0 nm corresponds to not performing any grouping. Predictions within a group are assigned a common set of emitter coordinates which is calculated as weighted average of their individual coordinates. We compare three different options for the weighted average: Uniform weighting (‘None’, solid lines); Weighting by the inferred number of photons for CSpline and DECODE or the inferred confidence for DeepSTORM3D (‘photons’, dotted line); Weighting by the predicted DECODE *σ* values, where the *x, y* and *z* values are individually weighted by 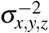. **a, b)**: 3D efficiencies across grouping radii. Grouping is especially useful in the low density setting (a) where DECODE without temporal context (DECODE single) with a correctly set grouping radius can match the performance of DECODE with temporal context (DEOCDE multi) without grouping. This is, however, only the case when weighting by the uncertainty estimates that DECODE provides. Using grouping on top of DECODE multi offers little additional benefit. **c, d)**: Number of groups divided by the number of localizations. Detecting all emitters and correctly grouping them would result in a ratio of 1 : 3 as on average each emitter is visible in three consecutive frames. See methods and Supplementary Table S1 for additional details on training and evaluation.

**Extended Data Figure 3.**
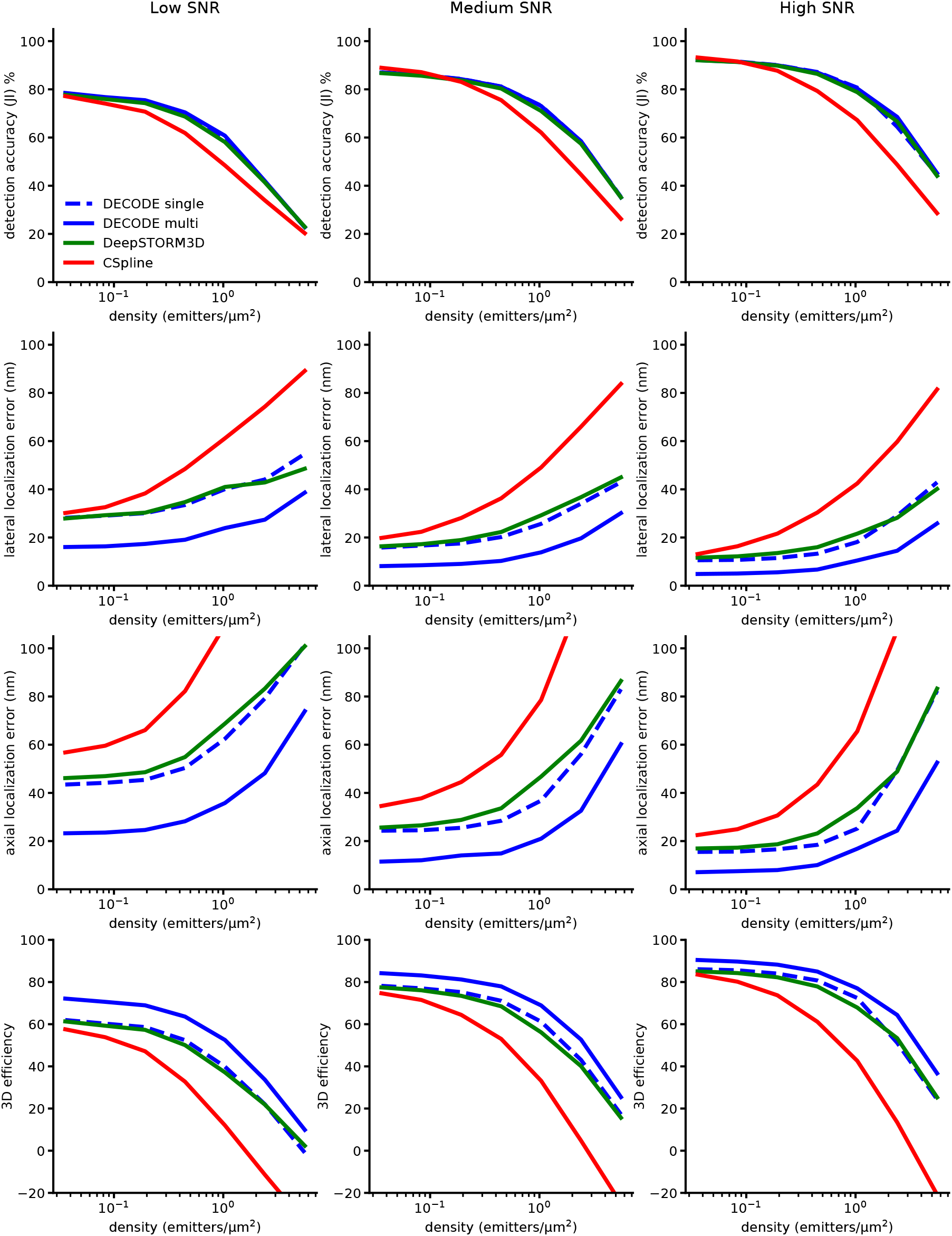
Comparison of performance metrics across densities and SNRs. DECODE outperforms DeepSTORM3D and CSpline across densities and SNRs. See methods and Supplementary Table S1 for additional details on training and evaluation.

**Extended Data Figure 4.**
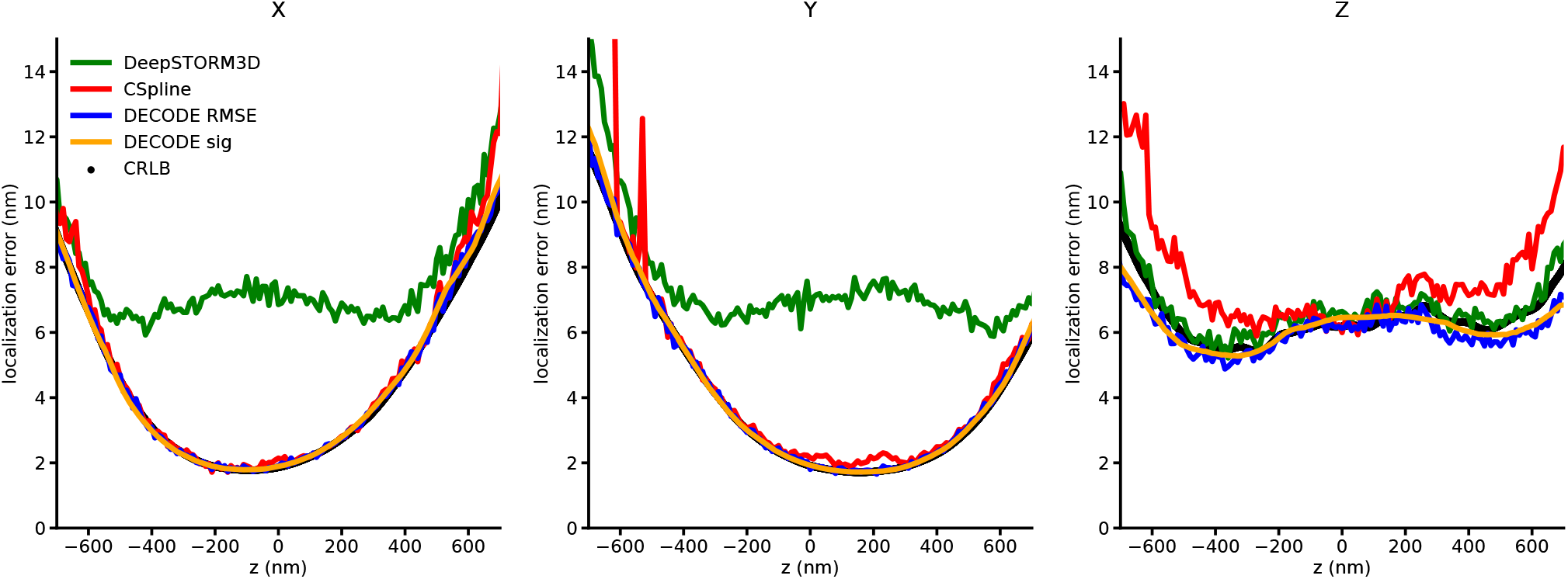
Comparison of localization error and CRLB for single emitter fitting. The RMSE achieved by DECODE and its predicted *σ* values closely match the single emitter CRLB in every dimension. CSpline is also able to achieve the CRLB, which has been shown for iterative MLE fitters before. In contrast the resolution that DeepSTORM3D can achieve is limited by its output representation and the size of the super-resolution voxels. RMSE and DECODE *σ* averaged over 10 nm bins. See methods and Supplementary Table S1 for additional details on training and evaluation.

**Extended Data Figure 5.**
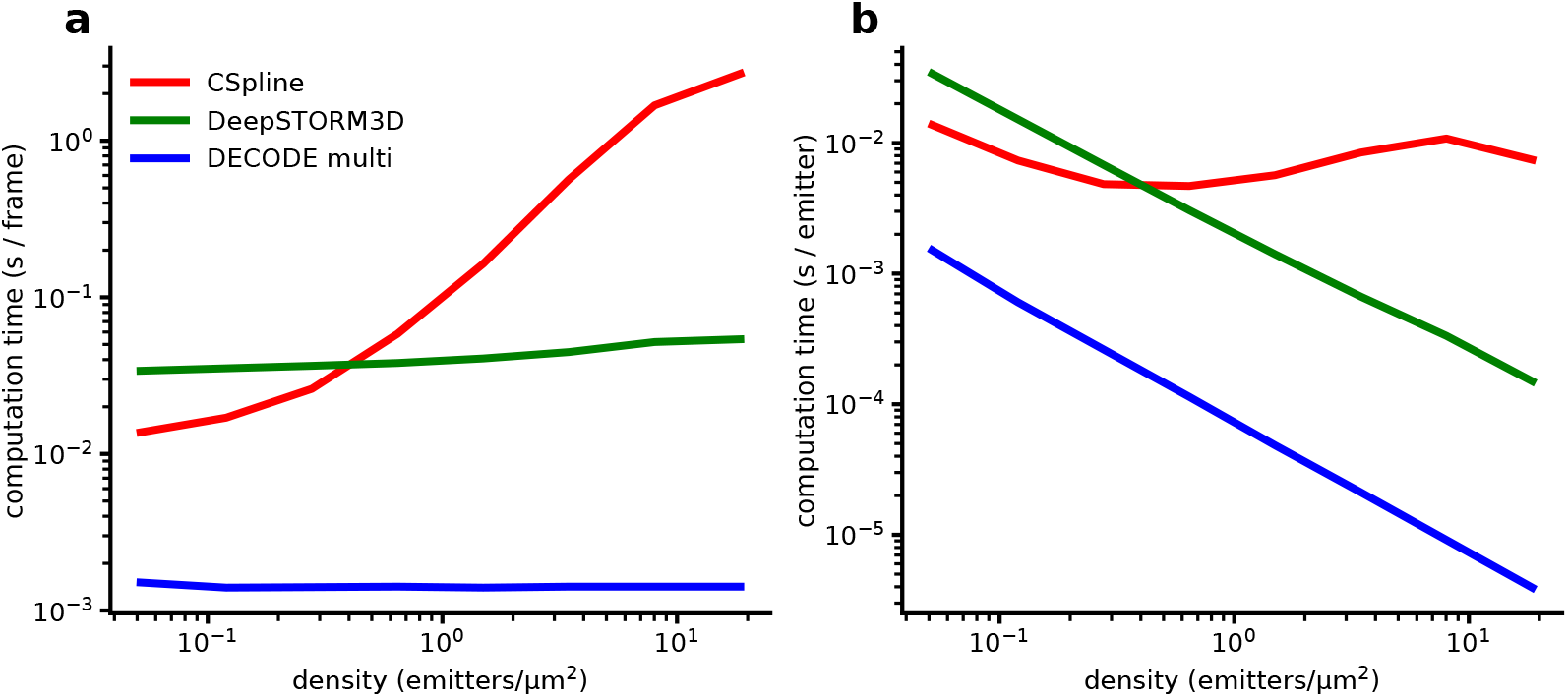
Comparison of computation times. **a)**. Measured as the time it takes to analyze a 64 × 64 pixel frame with varying emitter densities. Trained DECODE and DeepSTORM3D models were evaluated using a NVIDIA RTX2080Ti GPU. Computation time includes the network forward pass and post-processing and does not include training time. CSpline was evaluated on an Intel(R)Xeon(R) CPU E5-2697 v3. **b)** Computation time per simulated emitter. The computation time of CSpline scales with the number emitters while the two deep learning based approaches scale with the number (and size) of the analyzed frames. GPU-based DECODE is about 20 times faster than GPU-based DeepSTORM3D and outperforms CPU-based CSpline even at low densities.

**Extended Data Figure 6.**
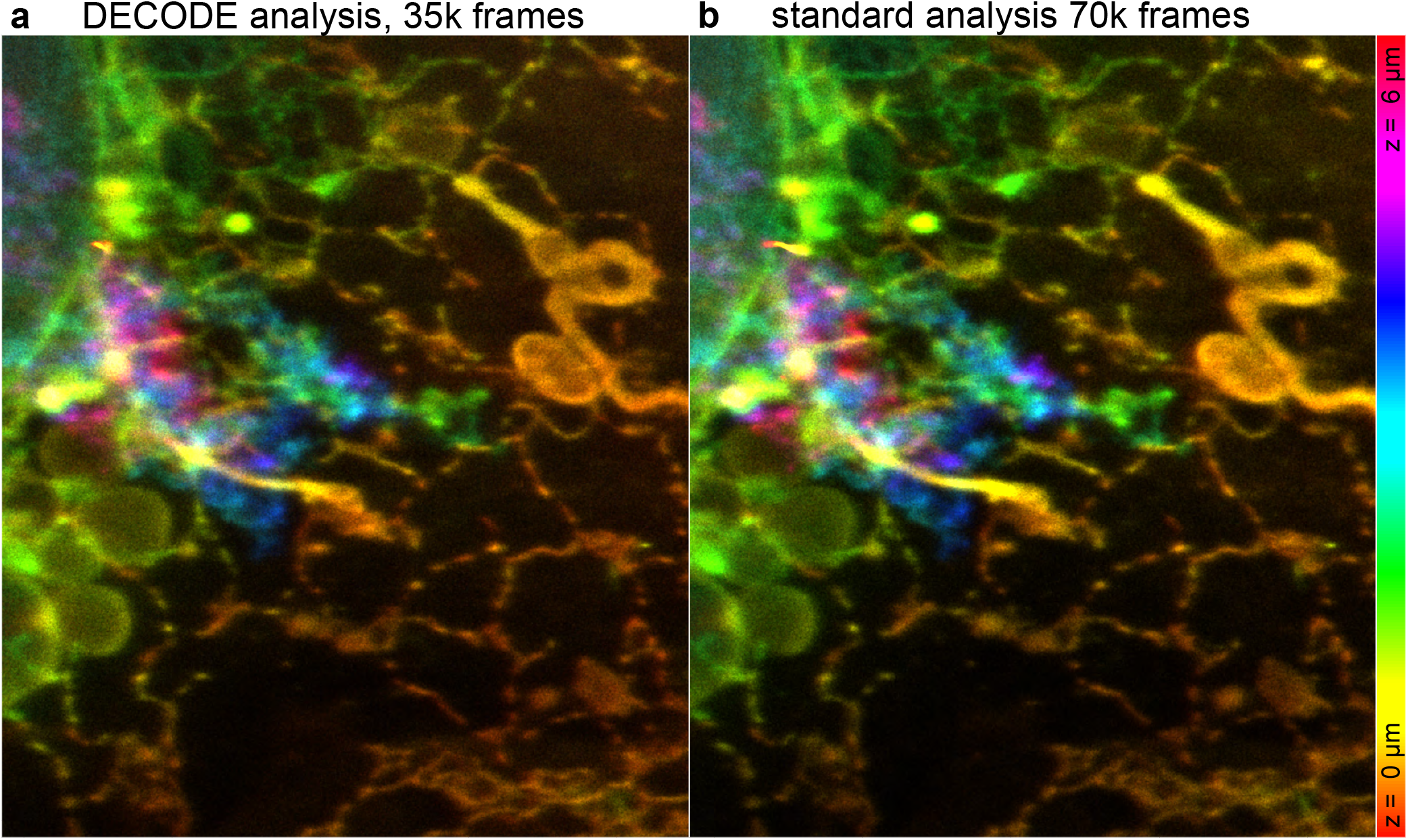
DECODE reduces acquisition times in LLS-PAINT. DECODE reconstruction of 35,000 frames (**a**) results in the same number of localizations as the Standard reconstruction of 70,000 frames (**b:**). As DECODE detects twice as many localizations as the traditional analysis, it needs only approx. half of the frames for a high-quality reconstruction.

**Extended Data Figure 7.**
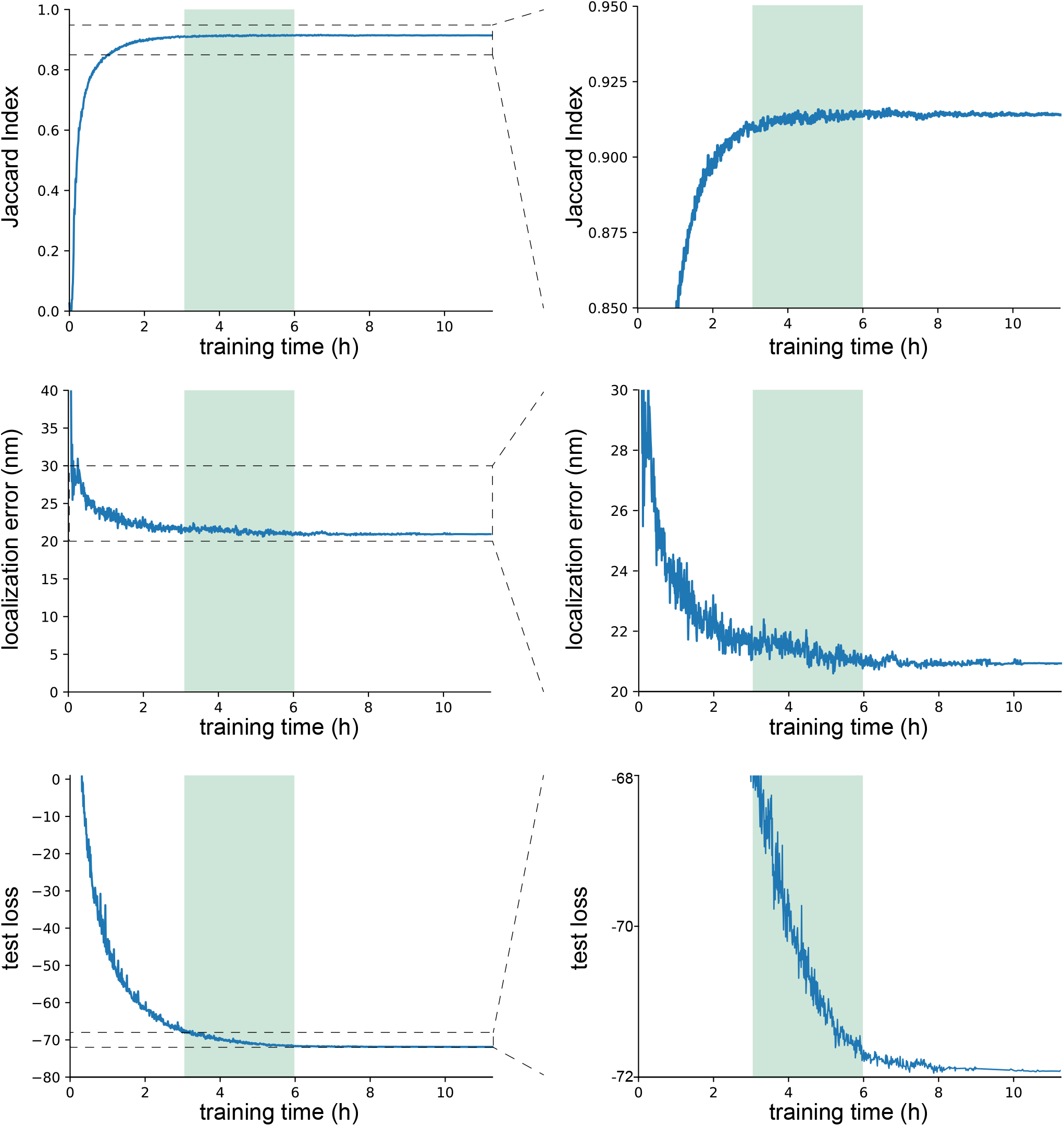
Performance as a function of deep network training time. Convergence of the accuracy of DECODE for several performance metrics. Runtimes are measured on a single nVidia RTX 2080 Ti GPU. The estimated training achievable with the maximum of 12 hours possible on the free tier of Google Colab is shown in green range (assuming that a Google Colab GPU is 2x-4x slower than the nVidia RTX 2080 Ti GPU). This suggests that acceptable performance is achievable using DECODE and Google Colab at minimal cost, no GPU needed. Metrics evaluated for prediction > 0.5 detection probability estimate without sigma filtering. Training data was simulated at high SNR (as described in Figure 2c) at an average density of 1 μm^−2^.

**Extended Data Figure 8.**
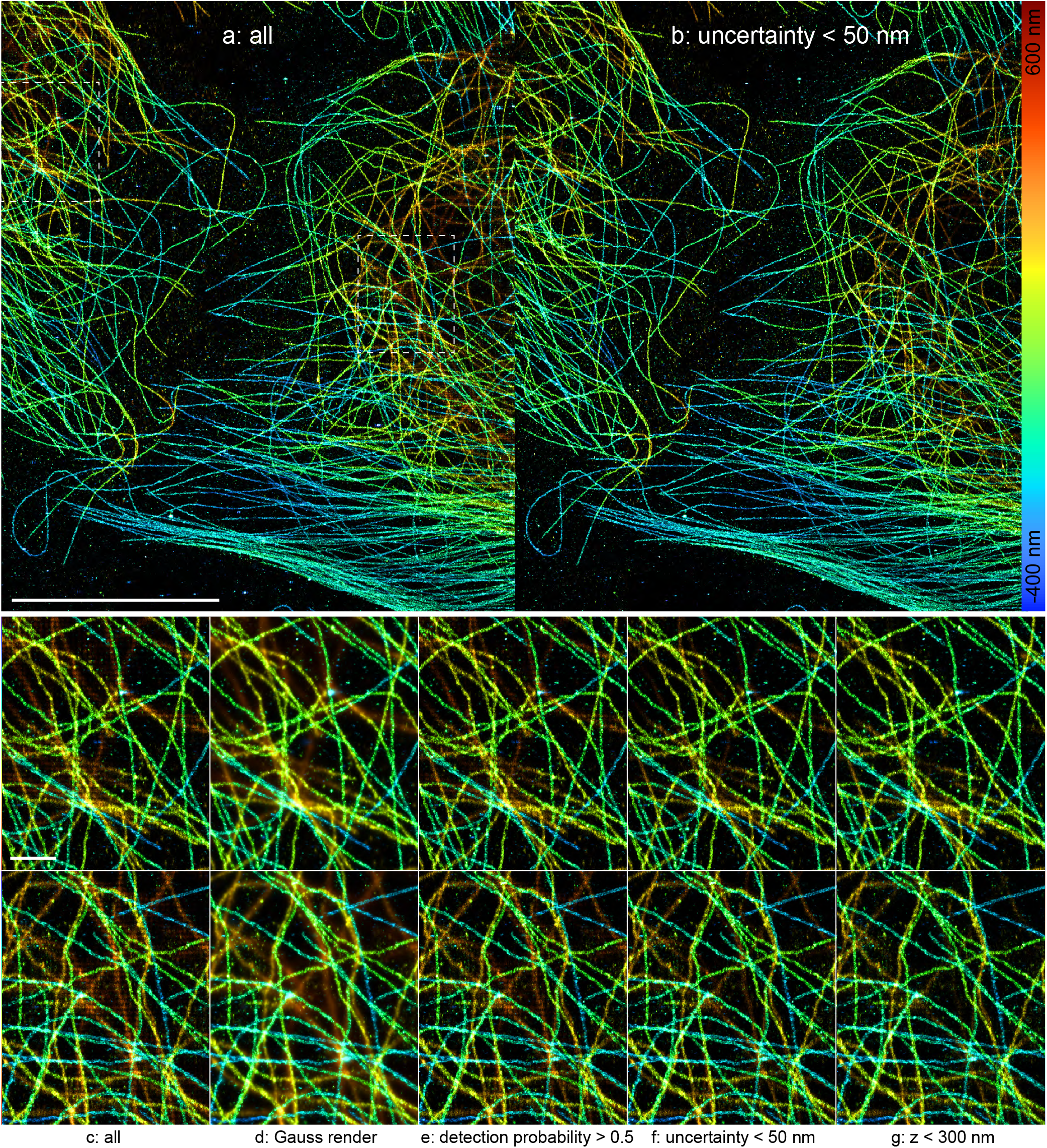
Removing Pixelation artifacts. Dim, dense out-of-focus localizations have a bias towards the pixel center (a,c). This bias is not visible if every localization is rendered as a Gaussian with a standard deviation equal to the predicted uncertainty *σ* (d). Filtering according to the detection probability reduces the artifact (e). Filtering according to the predicted uncertainty *σ* (b,f) or the fluorophore z-position (g) also removes the pixelation artifact. Scale bars 10 μm (a,b) and 1 μm (c-g).

## Supplementary Information

**Supplementary Figure S1.**
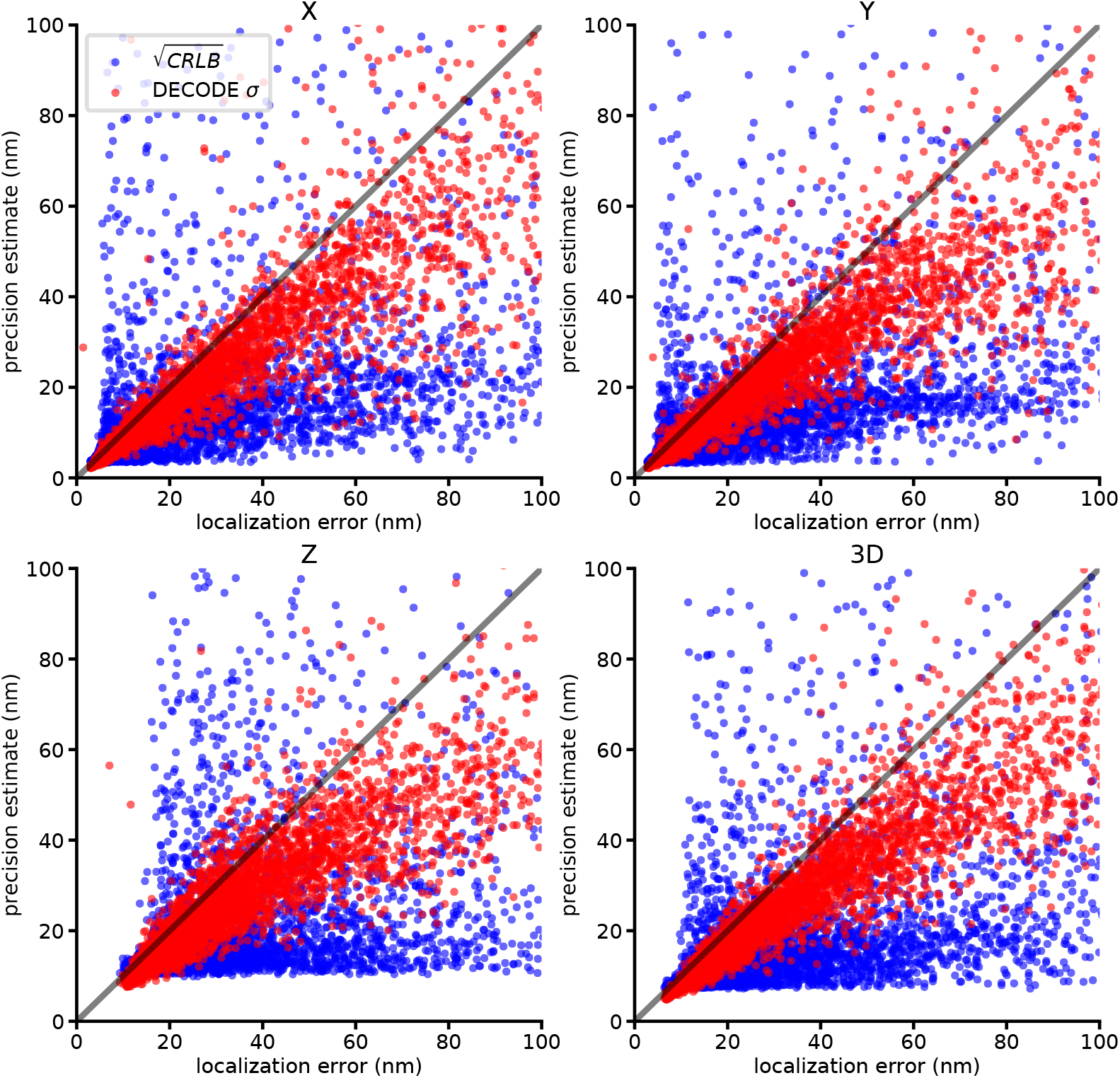
Comparison of inferred uncertainty and single emitter CRLB in each dimension for dense data. DECODE’s *σ* predictions correlate closely to the measured localization error in each dimension, i.e. much better than the single emitter CRLB estimate which assumes isolated emitters. We simulated the same dense emitter configuration 100 times and calculated the measured localization error as the RMSE of the predictions of the coordinates. See Methods and Supplementary Table S1 for additional details on training and evaluation.

**Supplementary Figure S2.**
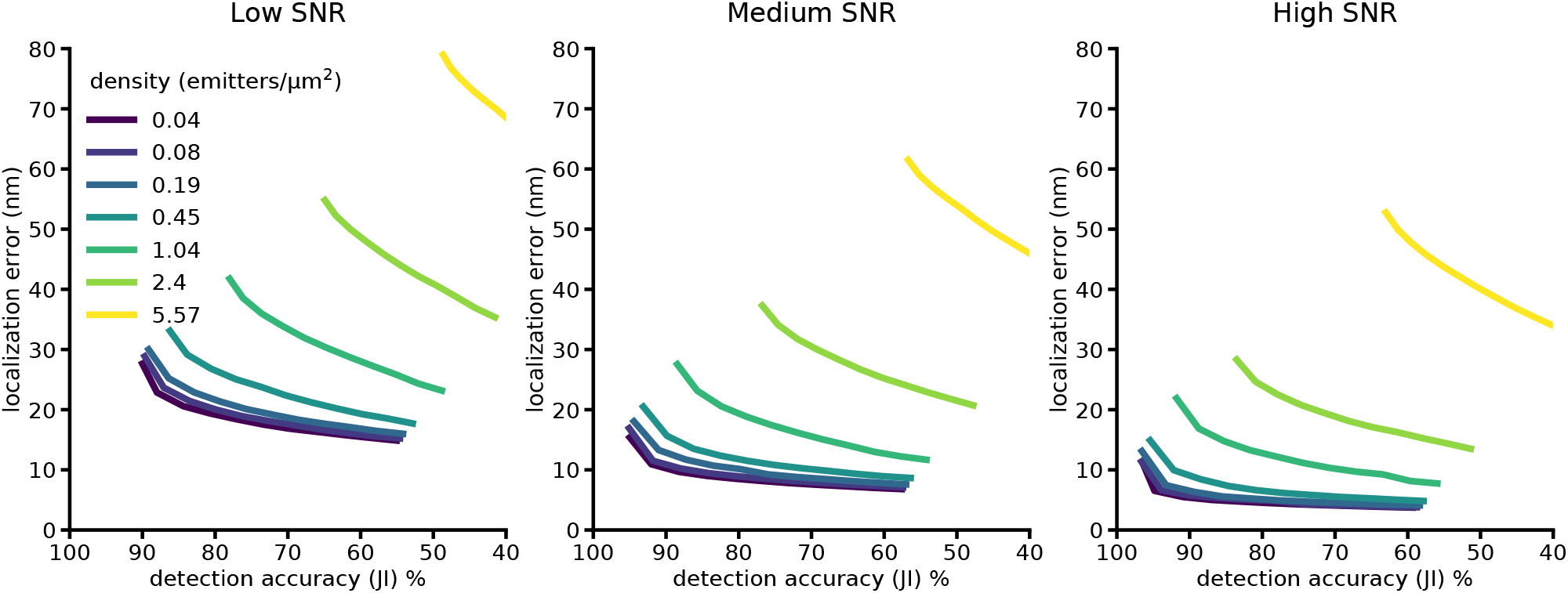
Impact of filtering on localization error and detection efficiency. Each line corresponds to the two performance metrics evaluated for DECODE multi predictions for a given density with 0% - 40% of the worst predictions removed (ordered by the predicted DECODE *σ*). As the DECODE *σ* are well calibrated they allow to effectively trade off detection accuracy for a lower localization error. See methods and Supplementary Table S1 for additional details on training and evaluation.

**Supplementary Figure S3.**
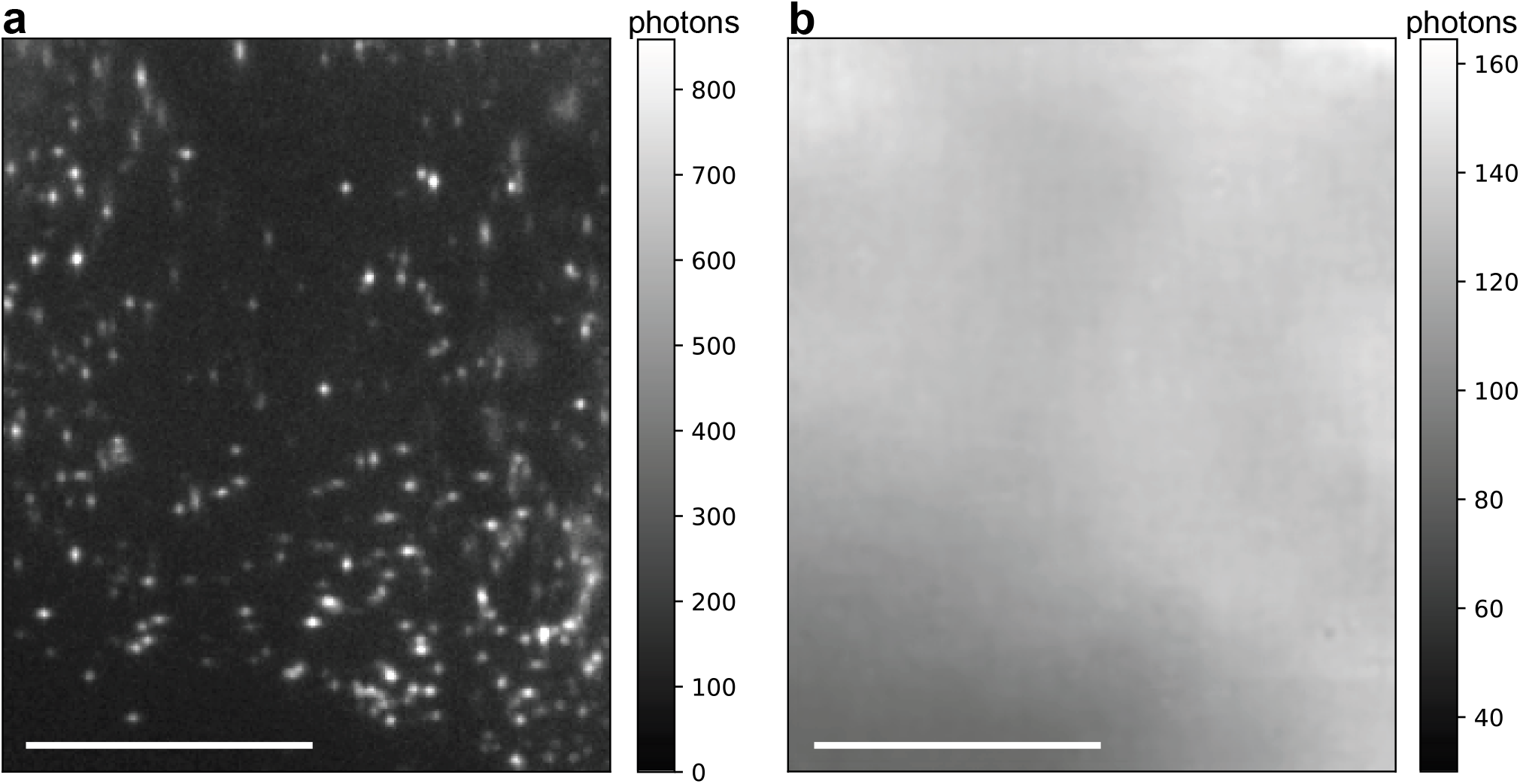
Background prediction. Scale bar 100 pixels. Depicted are raw input frame from the data set corresponding to Fig. 4e (converted into photon units) and respective background prediction output of the model (rescaled into photon units).

**Table S1.**
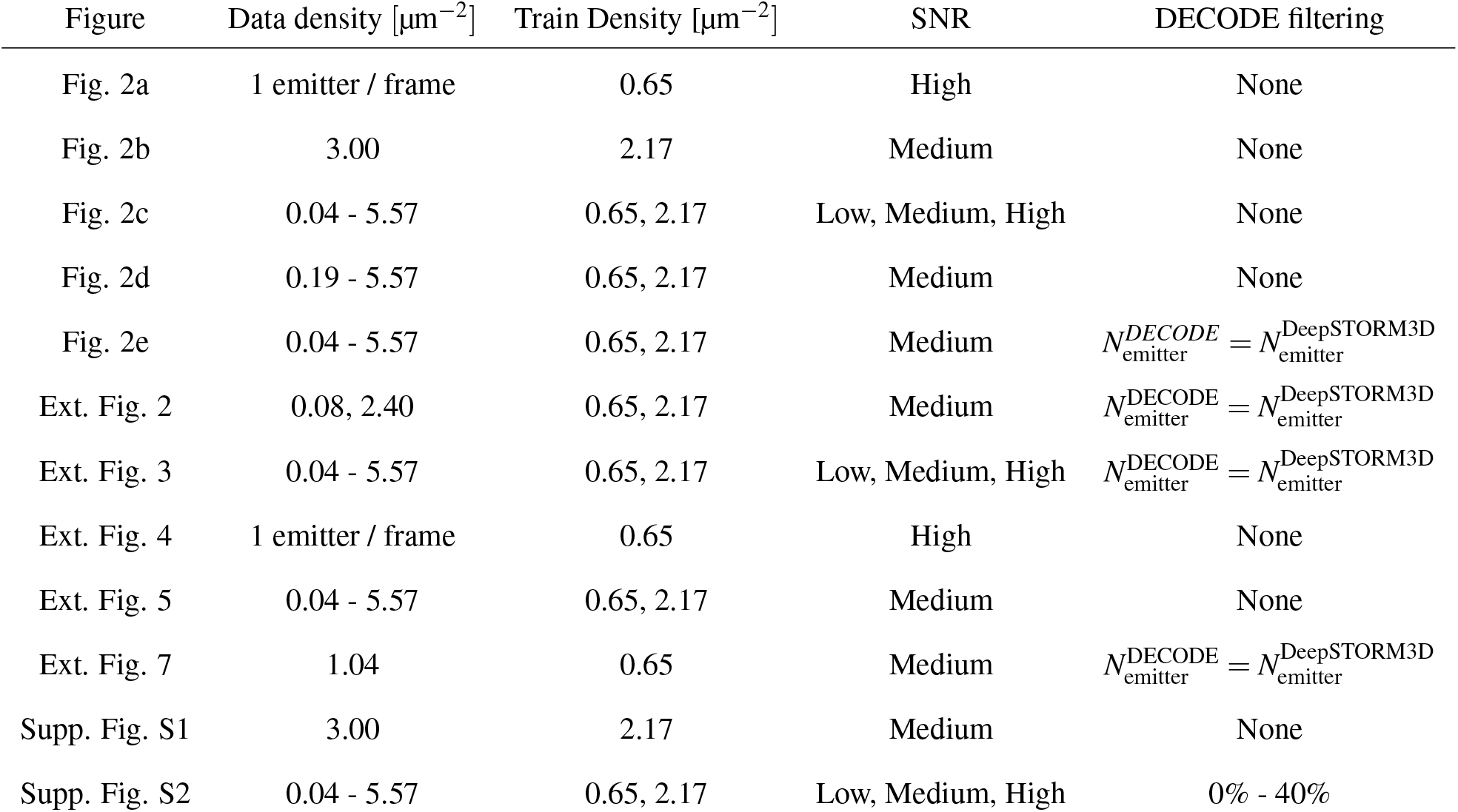
Simulation and evaluation parameters for experiments based on our own simulations. Data density refers to the set of frames used for evaluation, while train density is the density of the simulated frames used for training the DECODE and DeepSTORM3D networks. For each emitter, we draw a photon flux from a Gaussian distribution *N*(*μ*_flux_, *σ*_flux_). Low, medium and high SNR refer to mean photon counts of 1000, 5000 and 20,000 with background levels of 10, 50 and 200 photons per pixel respectively. The standard deviation of the intensity is calculated as *σ*_flux_ = *μ*_flux_/20. We used a mean on-time of 2 frames. For the CRLB comparisons in Fig. 2a and Extended Data Fig. 4 all emitters were instead simulated with exactly 20,000 photons and 200 background photons and did not persist across frames. To compare the DECODE’s *σ* predictions with the measured localization uncertainty (Fig. 2b and Supp. Fig. S1) we simulated 100 frames and sampled the noise 100 times. For the CRLB comparisons (Fig. 2a and Ext. Fig. 4) we simulated 100k frames. For the remaining figures we simulated frames for each combination of data density and SNR until we acquired at least 20k emitters and 1k frames.

## Footnotes

† http://bigwww.epfl.ch/smlm/challenge2016/leaderboard.html

